# Single-cell multi-omic analysis of thymocyte development reveals drivers of CD4/CD8 lineage commitment

**DOI:** 10.1101/2021.07.12.452119

**Authors:** Zoë Steier, Dominik A. Aylard, Laura L. McIntyre, Isabel Baldwin, Esther Jeong Yoon Kim, Lydia K. Lutes, Can Ergen, Tse-Shun Huang, Ellen A. Robey, Nir Yosef, Aaron Streets

## Abstract

The development of CD4 and CD8 T cells in the thymus is critical to adaptive immunity and is widely studied as a model of lineage commitment. Recognition of self-MHCI/II by the T cell antigen receptor (TCR) determines the lineage choice, but how distinct TCR signals drive transcriptional programs of lineage commitment remains largely unknown. We applied CITE-seq to measure RNA and surface proteins in thymocytes from wild-type and lineage-restricted mice to generate a comprehensive timeline of cell state for each lineage. These analyses revealed a sequential process whereby all thymocytes initiate CD4 lineage differentiation during an initial wave of TCR signaling, followed by a second TCR signaling wave that coincides with CD8 lineage specification. CITE-seq and pharmaceutical inhibition experiments implicate a TCR/calcineurin/NFAT/GATA3 axis in driving the CD4 fate. Overall, our data suggest that multiple redundant mechanisms contribute to the accuracy and efficiency of the lineage choice.

## Introduction

The commitment of a developing thymocyte into the CD4+ helper or CD8+ cytotoxic T cell fate provides an important model for understanding the interplay between signal transduction pathways and cell fate decisions and is one of the most intensely studied examples of a mammalian cell fate decision. The ultimate fate of a thymocyte is determined by the specificity of its T cell receptor (TCR) for major histocompatibility complex (MHC) molecules during positive selection in the thymus, with recognition of class I MHC (MHCI) leading to the CD8 fate, and recognition of class II MHC (MHCII) leading to the CD4 fate. CD8 and CD4 are coreceptors that aid in the recognition of MHCI and MHCII respectively, and the dynamic expression pattern of coreceptors plays an important role in lineage commitment (Germain, 2002; Xiong and Bosselut, 2012, Singer et al., 2008, Shinzawa et al., 2022). The most prominent model of lineage commitment, termed “kinetic signaling”, focuses on CD8 down-regulation in CD8-fated cells and the associated drop in TCR signaling (Singer et al., 2008). However, there is evidence that TCR signaling impacts CD4 and CD8 T cell development throughout the >2-day process of positive selection (Kisielow and Miazek, 1995; Liu and Bosselut, 2004; Au-Yeung et al., 2014; Sinclair and Seddon, 2014), and we do not have a clear picture of the quantitative and temporal pattern of TCR signaling throughout lineage specification. It is also not known whether different transcriptional targets of the TCR pathway are activated in a temporal and/or lineage specific manner. Thus, the molecular links between TCR signaling and induction of the lineage-defining transcription factors THPOK (encoded by *Zbtb7b*) and RUNX3 in mature CD4 and CD8 T cells respectively (Taniuchi, 2016) remain unknown.

A major complicating factor in addressing these questions is the large diversity in TCR specificity and resulting cell fates, with many cells fated to undergo death by neglect, negative selection, or agonist selection, alongside CD4-fated and CD8-fated cells. While the use of transgenic mice bearing fixed rearranged TCRs that mediate positive selection and a largely predetermined lineage choice can help address this challenge, even with a fixed TCR, defining cell states and ordering them into a developmental trajectory remains a major challenge. Traditionally, cell states have been characterized using flow cytometry with a limited number of surface protein markers including CD4 and CD8 (Germain, 2002). While the distinction between immature double positive (DP; CD4+CD8+) versus mature single positive (SP; CD4+CD8- and CD4-CD8+) can be relatively clear cut, defining “transitional” populations using intermediate levels of CD4 and CD8, together with other markers can be subjective, and the gates used to define transitional cells are often contaminated by early or later populations, as well as cells not fated for either the CD4 or CD8 lineage. This approach also lacks the ability to make quantitative global comparisons in gene expression between cell states.

More recently, advances in single-cell RNA sequencing (scRNA-seq) technologies have enabled the unbiased observation of transcriptional heterogeneity in the mammalian thymus. scRNA-seq has been used to construct a census of cell states in the thymus and helped identify new thymic subsets, as well as shed light on the kinetics of TCR rearrangement prior to positive selection (Park et al., 2020), identify the early precursor populations that seed the thymus (Lavaert et al., 2020), characterize early progenitor commitment (Zhou et al., 2019) and examine the CD4 versus CD8 lineage choice (Chopp et al., 2020; Karimi et al., 2021). While these studies provided important insights, they did not have sufficient temporal resolution of the CD4 versus CD8 commitment process to connect TCR signaling events to the induction of THPOK, RUNX3 and the initiation of lineage specific transcriptional programs. As a case in point, there is evidence for a late CD8-fated CD4+CD8+ thymocyte population (Saini et al., 2010), but this population was not distinguished from earlier TCR signaled CD4+CD8+ populations that contain both CD4-fated and CD8-fated cells (Chopp et al., 2020; Karimi et al., 2021). In addition, the lack of protein information limited the ability to relate these studies to prior knowledge on the diversity of cell states, accumulated by the rich flow cytometry-based literature. Generating a high-resolution delineation of the differentiation process that provides connections to flow cytometry-based studies is needed to inform models of lineage commitment and to identify early drivers of lineage divergence.

To address these challenges, we leveraged the CITE-seq protocol (Stoeckius et al., 2017) to simultaneously measure the transcriptome and over 100 surface proteins in thousands of single thymocytes as they transition from DP precursor into mature CD4 or CD8 T cells. Using the totalVI algorithm (Gayoso et al., 2021), we jointly analyzed the paired RNA and protein measurements in order to build a comprehensive, high resolution, timeline of RNA and protein expression changes spanning the phases of positive selection and commitment in both lineages. The availability of protein information allowed us to relate our findings to the foundational protein-based literature and add to it by clarifying intermediate developmental stages (including the late CD8-fated CD4+CD8+) and better define these stages by both transcript and surface protein composition. Finally, our study design included thymi from both wild-type (WT) and lineage-restricted mice, which enabled a comparison between CD4-versus CD8-fated cells at early stages when they are otherwise difficult to distinguish.

Through these analyses, we were able to address key questions about how TCR signals regulate lineage commitment. First, we were able to pinpoint the stage at which lineage-specific gene expression differences first emerged, and, through comparative analysis of lineage-restricted cells, identify TCR signaling through calcineurin-NFAT as a putative driver of the CD4 fate. Second, our analysis revealed two distinct temporal waves of TCR signaling: an early wave that is more sustained in CD4-fated cells, and a later wave that is specific to CD8-fated cells and that overlaps with CD8 lineage specification. CITE-seq data suggested that all the major branches of the TCR signaling pathway, including calcineurin-NFAT and MEK-ERK, were active in the first TCR signaling wave, however calcineurin-NFAT was less prominent in the later CD8-specific TCR signaling wave, again leading us to hypothesize a role for NFAT in CD4 lineage commitment. To validate this hypothesis, we applied drug perturbations to an ex vivo thymic slice culture system. While MEK inhibition broadly inhibited the development of CD4 and CD8 T cells, calcineurin inhibition selectively blocked the earliest phase of CD4 SP development. This corresponded to a failure of DP thymocytes to fully induce GATA3, a transcription factor known to be upstream of the CD4 commitment factor THPOK.

More broadly, our analysis provides a new understanding overview of the relationship between the two programs. We find that CD8-fated cells initially undergo a parallel, but transient, CD4 transcriptional program, including induction of THPOK, implying that CD8-fated cells audition for the CD4 fate prior to undergoing CD8 lineage specification. We also show that CD8-fated cells can co-express low levels of *Zbtb7b* and *Runx3* during the CD4 audition phase, implying that mutual antagonism between the two transcription factors is most active as CD8-fated cells transition from a failed CD4 audition to undergoing CD8 lineage specification.

By establishing a high-resolution map of development based on gene expression and surface proteins, our findings help fill the knowledge gap between TCR signaling at the cell surface and the subsequent differential activation of lineage controlling transcription factors in the nucleus. It also offers insight into the shared and unique components between the two programs, thus establishing a more comprehensive model for CD4 versus CD8 T cell fate commitment.

## Results

### A joint transcriptomic and surface protein atlas of thymocyte development in wild-type and lineage-restricted mice

To study T cell development and CD4/CD8 lineage commitment, we profiled thymocytes from both WT and lineage-restricted mice. Thymocyte populations in WT mice closely resemble those in humans (Park et al., 2020), and serve as a model of T cell development in a healthy mammalian system. However, in a WT system it is not possible to predict the ultimate fate of immature thymocytes, making it challenging to investigate the process of commitment to the CD4 or CD8 lineage. To probe the mechanism of lineage commitment, we profiled thymocytes from both WT (C57BL/6; referred to as B6) mice and mice with CD4-lineage- or CD8-lineage-restricted T cells. To track commitment to the CD4 lineage (MHCII-specific) we used mice that lack MHCI expression (β2M-/-; referred to as MHCI-/-), which have polyclonal TCR repertoires, and two TCR transgenic (TCRtg) mice that express TCRs that are specific for MHCII (AND and OT-II). To track commitment to the CD8 lineage (MHCI-specific) we used mice that lack MHCII expression (I-Aβ-/-; referred to as MHCII-/-) and two MHCI-specific TCRtg mice (F5 and OT-I). In all of these lineage-restricted mice, thymocytes are expected to pass through the same stages of development as WT thymocytes (Figure 1A). However, unlike in WT thymocytes, the fate of TCR-signaled thymocytes from lineage-restricted mice is known before cells present the CD4+ SP or CD8+ SP phenotype, allowing for independent characterizations of CD4 and CD8 T cell development and lineage commitment.

**Figure 1:**
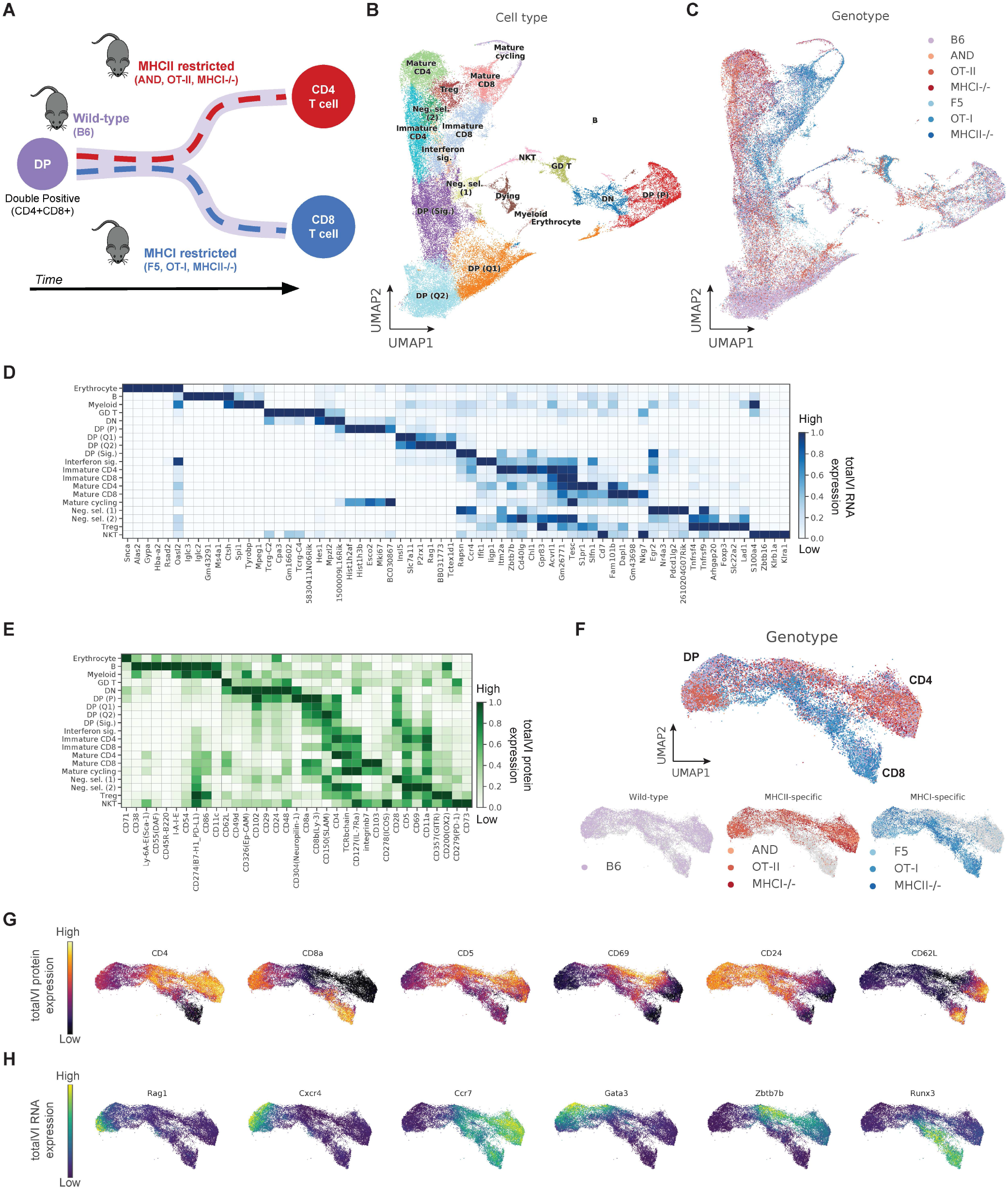
A joint transcriptomic and surface protein atlas of thymocyte development in wild-type and lineage-restricted mice. **(A)** Schematic representation of thymocyte developmental trajectories in WT, MHCII-restricted (MHCI-/-, OT-II TCRtg and AND TCRtg), and MHCI-restricted (MHCII-/-, OT-I TCRtg, and F5 TCRtg) mice used in CITE-seq experiments. **(B, C)** UMAP plots of the totalVI latent space from all thymocyte CITE-seq data labeled by **(B)** cell type annotation and **(C)** mouse genotype. **(D, E)** Heatmaps of markers derived from totalVI one-vs-all differential expression test between cell types for **(D)** RNA and **(E)** proteins. Values are totalVI denoised expression. **(F)** UMAP plots of the totalVI latent space from positively-selected thymocytes with cells labeled by mouse genotype. **(G, H)** UMAP plots of the totalVI latent space from positively-selected thymocytes. **(G)** Cells colored by totalVI denoised expression of protein markers of lineage (CD4, CD8a), TCR signaling (CD5, CD69), and maturation (CD24, CD62L). **(H)** Cells colored by totalVI denoised expression of RNA markers of TCR recombination (*Rag1*), thymic location (chemokine receptors *Cxcr4*, *Ccr7*), and lineage regulation (transcription factors *Gata3*, *Zbtb7b*, *Runx3*). GD T, gamma-delta T; DN, double negative; DP (P), double positive proliferating; DP (Q1), DP quiescent 1; DP (Q2), DP quiescent 2; DP (Sig.), DP signaled; Interferon sig., interferon signature; Neg. sel. (1), negative selection wave 1; Neg. sel. (2), negative selection wave 2; Treg, regulatory CD4 T cell; NKT, natural killer T.

We characterized thymocyte development at the single-cell level by measuring transcriptomes and surface protein composition using CITE-seq (Stoeckius et al., 2017) with a panel of 111 antibodies (Table S1). We jointly analyzed these features using totalVI, which accounts for nuisance factors such as variability between samples, limited sensitivity in the RNA data and background in the protein data (Gayoso et al., 2021; Methods). We collected thymi from two biological replicates per lineage-restricted genotype and five WT biological replicates (Table S2). Because the majority of thymocytes from TCRtg mice undergo positive selection, we analyzed TCRtg thymocytes without enrichment, while samples from non-transgenic mice, MHC-deficient samples and three WT replicates, were sorted by FACS for CD5+TCRβ+ to enrich for thymocytes undergoing positive selection (Figure S1A). We integrated CITE-seq data from all samples (72,042 cells) using totalVI, which allowed us to stratify cell types and states based on both RNA and protein information regardless of mouse genotype (Figure 1B). We identified the expected stages of thymocyte development including early CD4-CD8- (double negative; DN) and proliferating CD4+CD8+ (double positive proliferating; DP (P)) stages. We detected non-proliferating preselection stages of DP cells (double positive quiescent 1 and quiescent 2; DP (Q1, Q2)) undergoing TCR recombination, as well as DP cells post-recombination that are downregulating *Rag* and receiving positive selection signals (signaled DP). In addition to immature and mature stages of CD4 and CD8 T cells, we observed two distinct waves of cells that appeared to be undergoing negative selection based on expression of *Bcl2l11* (BIM), *Nr4a1* (NUR77), and *Ik2f2* (Helios) (Daley et al., 2013) (Figure S1B). The first wave appeared to emerge from the signaled DP population (lying adjacent to a cluster of dying cells, and the second wave emerged from immature CD4 T cells. The two negative selecting clusters also express markers described by (Daley et al., 2013) that distinguish the early and late waves of negative selection. These include upregulation of *Pdcd1* (PD-1) (Figure 1E) and downregulation of CD4 and CD8 mRNA (Figure S1B) and protein (Figure S1C) by the Negative selection (1) cluster, and upregulation of *Tnfrsf18* (CD357/GITR), *Tnfrsf4* (CD134/OX40) by Negative selection (2) cluster (Figure S1B-C). *Foxp3*+ regulatory T cells appeared to cluster near mature conventional CD4 T cells and the second subset of negatively selected cells. Other populations included unconventional T cells (gamma-delta T cells, NKT cells), small clusters of non-T cells (B cells, myeloid cells, and erythrocytes), a thymocyte population with high expression of interferon response genes (Xing et al., 2016), and a population of mature T cells that had returned to cycling following the cell cycle pause during thymocyte development. As expected, WT, MHCII-specific, and MHCI-specific samples were well-mixed in earlier developmental stages but branched into CD4 and CD8 lineages in later-stage populations (Figure 1C).

Using totalVI, we defined cell populations with traditional cell type markers (Figure S1B-C) and with unbiased differential expression tests of all measured genes and proteins (Figure 1D-E, Table S3). Top differentially expressed features included classical cell surface markers of lineage (e.g., CD4, CD8), key transcription factors (e.g., *Foxp3*, *Zbtb7b*), and markers of maturation stage (e.g., *Rag1*, *Ccr4*, *S1pr1*). In addition to supporting the relevance of surface proteins in characterizing cell identities, these multi-omic definitions revealed continuous expression changes, particularly between the DP and CD4 and CD8 SP stages, that are best understood not as discrete populations, but as part of a continuous developmental process. Observation of these groups allowed us to select the populations of thymocytes receiving positive selection signals for further continuous analysis.

We focused our analysis on developing thymocytes from the signaled DP stage through mature CD4 and CD8 T cells (Methods; Figure S1D). The totalVI latent space derived from these populations captured the continuous transitions that stratified thymocytes by developmental stage and CD4/CD8 lineage (Figure 1F). This was evident through visualization of totalVI denoised protein and RNA expression of known markers including, for example, CD4 and CD8 protein expression, which revealed a transition into a CD4 or CD8 SP phenotype in each of the two branches. In the CD8 branch, a CD4+CD8+ population could be seen even after a separation of the lineages in the UMAP representation of the totalVI latent space (Figure 1G; Becht et al., 2019), indicating that combining transcriptome-wide information with surface protein measurements might reveal earlier signs of lineage commitment than could have been observed from FACS-sorted populations. We also observed protein markers indicative of positive-selection-induced TCR signaling (CD5, CD69) and maturation stage (CD24, CD62L), as well as RNA markers of TCR recombination (*Rag1)*, cell location within the thymus (chemokine receptors *Cxcr4*, *Ccr7*), and lineage regulation (transcription factors *Gata3*, *Zbtb7b*, *Runx3*) (Figure 1H). This single-cell analysis of positively-selecting thymocytes from WT and lineage-restricted mice thus provides a high-resolution snapshot of the continuous developmental processes, capturing cells at a variety of states that span the spectrum between the precursor (DP) and late (CD4 or CD8 SP) stages.

### Pseudotime inference captures continuous maturation trajectory and clarifies intermediate stages of development

To further characterize the observed continuum of cell states, we performed pseudotime inference with Slingshot (Street et al., 2018; Saelens et al., 2019) to delineate the changes in RNA and protein expression over the course of development from the DP to SP stages (Figures 2A and S2A; Methods). Since cell-cell similarities in the reduced dimension space were based on both RNA and protein information, the placement of a cell in pseudotime reflected continuous changes in both gene and protein expression. The pseudotime inferred by Slingshot consisted of a branching trajectory with each branch corresponding to a different lineage (CD4, CD8), as can be observed by the positioning of cells from lineage-restricted mice (Methods; Table S4 and Figure 1F).

We confirmed that the pseudotime ordering determined by Slingshot correctly captured the presence and timing of many known expression changes during thymocyte development (Hogquist et al., 2015) at the RNA and protein levels (Figure 2B-C). This included events such as early downregulation of TCR recombination markers *Rag1* and *Rag2*, continuous downregulation of early markers such as *Ccr9* and *Cd24a*/CD24, transient expression of the activation and positive selection marker *Cd69*/CD69, and late upregulation of maturation markers such as *Klf2*, *S1pr1*, and *Sell*/CD62L. To explore beyond known markers, we tested for differential expression over pseudotime with totalVI (Methods) and created a comprehensive timeline of changes in RNA and protein expression separately for each lineage (Figure 2D; Tables S5 and S6). There were some expected differences visible between the two lineages in the expression of key molecules (e.g., coreceptors and master regulators; Figure 2B). However, known markers of maturation followed similar patterns and timing in both lineages (Figure 2C). Furthermore, some of the most significant differential expression events over time were common between the two lineages including the early downregulation of *Arpp21* (a negative regulator of TCR signaling (Mingueneau et al., 2013)), transient expression of the transcriptional regulator *Id2* (Cannarile et al., 2006), upregulation of Tesc, (an inhibitor of calcineurin (Perera et al., 2010)), and later upregulation of *Ms4a4b* (shown to inhibit proliferation in response to TCR stimulation (Yan et al., 2012)) (Figure 2D). This pseudotime analysis therefore provides a comprehensive view of the continuum of expression changes during the developmental process. Moreover, the consistency in timing of expression changes enables an investigation of the two lineages at comparable developmental stages.

**Figure 2:**
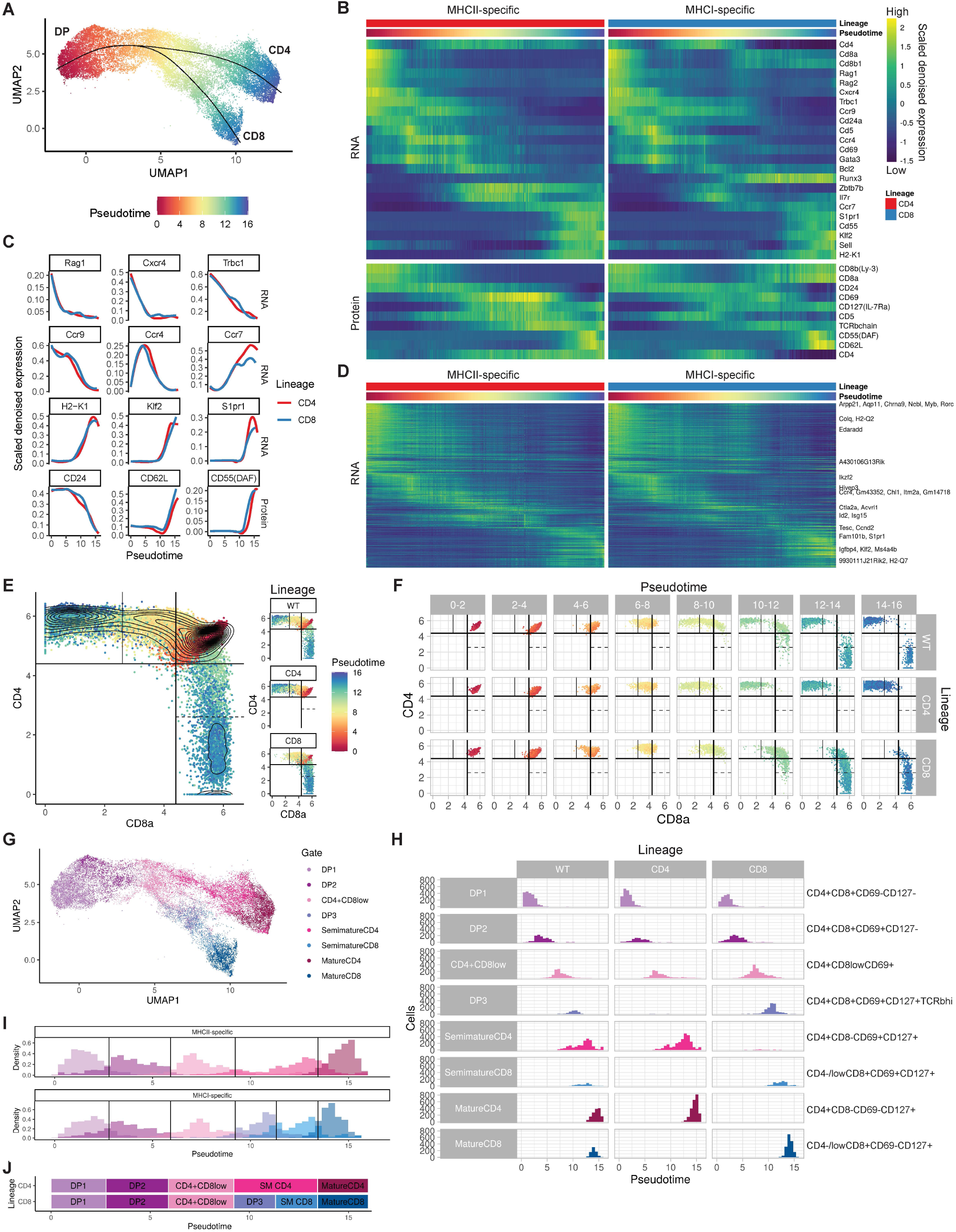
Pseudotime inference captures continuous maturation trajectory and clarifies intermediate thymocyte stages. **(A)** UMAP plot of the totalVI latent space from positively-selected thymocytes with cells colored by Slingshot pseudotime and smoothed curves representing the CD4 and CD8 lineages. **(B)** Heatmap of RNA (top) and protein (bottom) markers of thymocyte development over pseudotime in the CD4 and CD8 lineages. Features are colored by totalVI denoised expression, scaled per row, and sorted by peak expression in the CD4 lineage. Pseudotime axis is the same as in (A). **(C)** Expression of features in the CD4 and CD8 lineages that vary over pseudotime. Features are totalVI denoised expression values scaled per feature and smoothed by loess curves. **(D)** Heatmap of all RNA differentially expressed over pseudotime in any lineage. Features are scaled and ordered as in (B). Labeled genes are highly differentially expressed over time (Methods). **(E)** In silico flow cytometry plots of log(totalVI denoised expression) of CD8a and CD4 from positively-selected thymocytes (left) and the same cells separated by lineage (right). Cells are colored by pseudotime. Gates were determined based on contours of cell density. **(F)** In silico flow cytometry plot of data as in (E) separated by lineage and pseudotime. **(G)** UMAP plot of the totalVI latent space from positively-selected thymocytes with cells colored by gate. Cells were computationally grouped into eight gates using CD4, CD8a, CD69, CD127(IL-7Ra), and TCRβ. **(H)** Histograms of cells separated by lineage and gate with cells colored by gate as in (G). **(I)** Stacked histograms of gated populations in MHCII-specific (top) and MHCI-specific (bottom) thymocytes, with thresholds classifying gated populations over pseudotime (Methods). **(J)** Schematic timeline aligns pseudotime with gated populations, with population timing determined as in (I).

We next sought to use pseudotime information to clarify the intermediate stages of development in the two lineages. Thymocyte populations have been commonly defined by surface protein expression using flow cytometry, and various marker combinations and gating strategies have been employed to subset thymocyte populations based on maturity and lineage (Germain, 2002; Xiong and Bosselut, 2012; Saini et al., 2010; Hu et al., 2012). However, due to the continuous nature of developmental intermediates, as well as technical variations in marker detection, no uniform consensus has emerged on how to define positive selection intermediates by flow cytometry. To address this, we performed in silico flow cytometry analysis on totalVI denoised expression of key surface protein markers and explored their ability to distinguish between different pseudotime phases along the two lineages (Methods).

Starting with the canonical CD4 and CD8 markers, we found that MHCII-specific cells appeared to progress continuously in pseudotime from DP to CD4+CD8low to CD4+CD8- (Figures 2E-F and S2B). In contrast, MHCI-specific cells appeared to progress from DP to the CD4+CD8low gate before reversing course to reach the eventual CD4-CD8+ gate later in pseudotime, consistent with the previous literature (Lundberg et al., 1995; Lucas and Germain, 1996; Chan et al., 1993). Specifically, while at pseudotime 6-8 nearly all MHCI-specific cells fall in the CD4+CD8low gate, at pseudotime 8-12 MHCI-specific thymocytes pass again through a DP phase, while the MHCII-specific lineage does not contain late-time DP cells. Although such a population of later-time MHCI-specific DP cells has been previously described (“DP3”; Saini et al., 2010), it is not commonly accounted for (Park et al., 2020; Chopp et al., 2020, Karimi et al., 2021), resulting in a missing stage of CD8 T cell development and potential contamination of the DP gate with later-time CD8 lineage cells. Our continuous view of the development process therefore provides an opportunity to explore the progression of MHCI-specific cells after the CD4+CD8low gate, which has been less well defined in the literature.

We next sought to computationally identify a minimal set of surface markers for these and other stages of differentiation in our data. Our goal was twofold: first, we aimed to leverage markers that had previously been used for isolating intermediate subpopulations to establish consistency between our data and other studies; second, we aimed to better characterize the late MHCI-specific DP stage using a refined, data-driven, gating strategy. We found that four stages in time (independent of lineage) could be largely separated by in silico gating on CD69 and CD127(IL-7Ra), in which thymocytes begin with low expression of both markers, first upregulate CD69, later upregulate CD127, and finally downregulate CD69 (Figure S2C-D). The addition of CD4 and CD8 as markers allowed for the separation of lineages at later times. Finally, we established that the later-time DP population that is prominent in the CD8 lineage could be distinguished from the earlier DP cells by high expression of TCRβ (Saini et al., 2010; Marodon et al., 1994) in addition to expression of both CD69 and CD127 (Figure S2D-E). Here, we refer to this later-time DP population as DP3 to distinguish it from the earlier DP1 (CD69-, CD127-) and DP2 (CD69+, CD127-) populations.

In combination, a gating scheme based on these five surface proteins (CD4, CD8, TCRβ, CD69, and CD127) identified eight populations (DP1, DP2, CD4+CD8low, DP3, semimature CD4, semimature CD8, mature CD4, and mature CD8). This scheme, which included a marker previously unused in this setting (CD127), allows FACS to approximate the binning of thymocytes along pseudotime and lineage (Figure 2G-J). Fluorescence-based flow cytometry replicated these CITE-seq-derived gates (Methods), enabling the isolation of the eight described populations and supporting the presence of the proposed intermediate stages (Figure S2F-G). Collectively, these findings allowed us to specify an updated model of positive selection intermediates in both the CD4 and CD8 lineages (Figure S2F).

### Paired measurements of RNA and protein reveal the timing of major events in CD4/CD8 lineage commitment

While defining thymocyte populations by cell surface markers provides an approximate ordering of discrete developmental stages, a quantitative and high-resolution timeline of the differences between the lineages provided by CITE-seq data could address key outstanding questions. In particular, while it is known that both CD4-fated and CD8-fated cells initially downregulate CD8, it is not clear whether MHCII- and MHCI-specific thymocytes exhibit parallel temporal changes in CD4-defining transcription factors. In addition, the key events that lead to CD8 lineage specification have not been defined, due in large part to the lack of temporal resolution for the events that occur in MHCI-specific thymocytes after divergence from the CD4 lineage. To gain insight into CD4 versus CD8 lineage commitment, we used our pseudotime analysis to compare the continuum of expression changes of TCR signaling targets, key transcription factors, and coreceptors involved in this process.

Previous studies have implicated more prolonged TCR signaling as a driver of CD4 lineage commitment, whereas the role of TCR signals in CD8 SP development remains controversial (Germain, 2002; Xiong and Bosselut, 2012, Singer et al., 2008). When viewing the dynamics of expression changes over pseudotime, we generally observed an expected pattern by which RNA expression preceded the corresponding change in protein expression, likely due to the time needed for protein translation and transport. The induction and repression of *Cd69*/CD69 exemplifies this trend (Figure S3A). As expected, we observed a greater TCR response (as indicated by expression of TCR target genes *Cd69* and *Egr1*) early in the CD4 lineage (Figure 3A), which became significantly different by pseudotime bin 4-5, corresponding with the DP2 stage (Figure 3B). Surprisingly, this pattern reversed at later pseudotimes, with higher TCR signals in CD8-fated cells by pseudotime bin 9-10, corresponding to the DP3 stage. Strikingly, *Cd69* and *Egr1* expression over pseudotime reveal two distinct waves of TCR signaling during positive selection: an initial wave that is broader in CD4-fated cells and a later wave that is specific for CD8-fated cells (Figure 3A and S3B).

**Figure 3:**
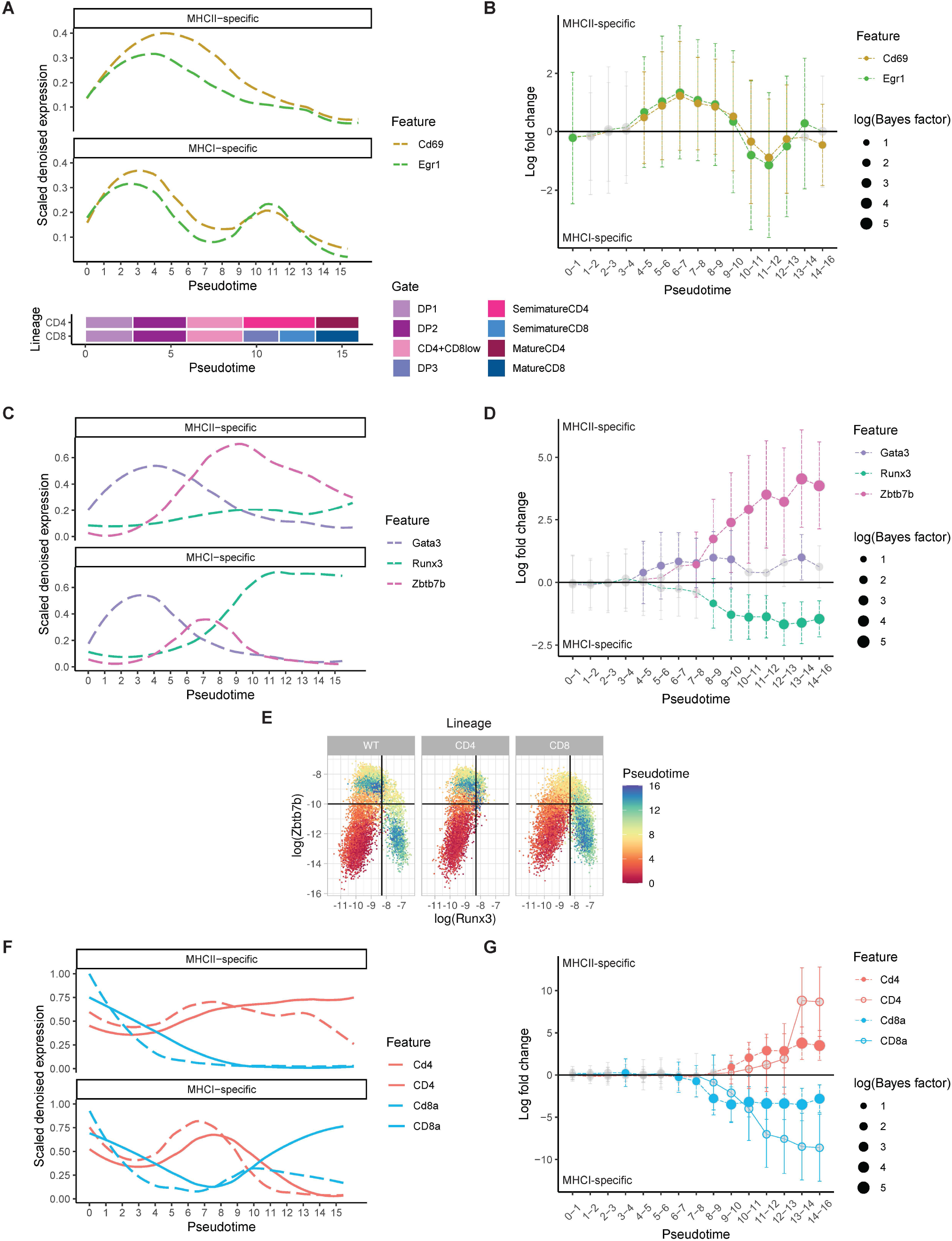
Paired measurements of RNA and protein reveal the timing of major events in CD4/CD8 lineage commitment. **(A)** Expression over pseudotime of TCR signaling response molecules. Features are totalVI denoised expression values scaled per feature and smoothed by loess curves. Schematic timeline below the plot aligns pseudotime with gated populations (see Figure 2J). **(B)** Differential expression over pseudotime between CD4 (MHCII-specific) and CD8 (MHCI-specific) lineages for RNA of TCR signaling response molecules. Non-significant differences are gray, significant differences are filled circles. Size of the circle indicates log(Bayes factor). Error bars indicate the totalVI-computed standard deviation of the median log fold change. **(C)** Expression over pseudotime as in (A) for RNA of key transcription factors. **(D)** Differential expression over pseudotime as in (B) for RNA of key transcription factors. **(E)** In silico flow cytometry plots of log(totalVI denoised expression) of *Runx3* and *Zbtb7b* from positively-selected thymocytes separated by lineage and colored by pseudotime. **(F)** Expression over pseudotime as in (A) for coreceptor RNA (dashed) and protein (solid). **(G)** Differential expression over pseudotime as in (B) for RNA of coreceptors. Significant RNA results are filled circles and significant protein results are open circles.

While previous studies characterized the expression of key lineage-defining transcription factors using flow cytometry (Muroi et al., 2008, Egawa and Littman, 2008), our pseudotime analysis could more precisely quantify differential expression of transcription factors as they diverge between the two lineages and relate this information directly to the timing of expression of other relevant genes. Of particular interest is the timing of TCR signaling relative to differential expression of the key lineage-specific transcription factor genes *Runx3* (CD8 lineage), *Zbtb7b* (encoding THPOK; CD4 lineage), and *Gata3* (a known up-stream activator of *Zbtb7b* that is more highly expressed in the CD4 lineage) (Wang et al., 2008, Taniuchi, 2016) (Figure 3C-D). We observed that the induction and differential expression of *Gata3* in CD4-fated cells coincided with differential expression of the first wave of TCR signaling genes in pseudotime bin 4-5 (corresponding to DP2), followed by differential expression of *Zbtb7b* in bin 7-8 (corresponding to CD4+CD8low stage). Differential upregulation of *Runx3* in CD8-fated cells occurred between pseudotimes 8-10, overlapping with DP3 and the second rise in TCR signaling in CD8-fated cells. Intracellular flow cytometry staining of thymocytes from wild-type mice supported the observed timing in differential expression of transcription factors (Figure S3C). While it has been suggested that IL7 and other STAT5 activating cytokines promote *Runx3* upregulation and the CD8 fate (Singer et al., 2008), we did not observe a lineage specific increase in STAT5 target gene expression correlating with *Runx3* upregulation (Figure S3D).

Examining continuous expression changes over pseudotime (Figures 3C and S3B), we observed a strikingly parallel pattern in both CD4 and CD8 lineage cells in which *Gata3* induction was followed by a rise in *Zbtb7b*. The expression of both CD4-associated transcription factors was lower and more transient in the CD8 lineage compared to the CD4 lineage, suggesting that both lineages initially “audition” for the CD4 fate, although MHCI-specific cells do so unsuccessfully. Interestingly, the large rise in *Runx3* expression, which occurred only in the CD8 lineage, overlapped with the decrease in *Zbtb7b* in that lineage (pseudotime 7-11, corresponding to the CD4+CD8low and DP3 stages). This implied that both transcription factors may be transiently co-expressed in CD8-fated cells, in spite of their ability to repress each other’s expression and their reported mutually exclusive expression at later stages (Egawa and Littman, 2008; Vacchio and Bosselut, 2016; Taniuchi, 2016). In silico flow cytometry of *Zbtb7b* and *Runx3* expression for WT or lineage-restricted thymocytes (Figures 3E and S4A) showed a continuous transition in which all thymocytes initially upregulated *Zbtb7b*, whereas CD8 lineage (MHCI-specific) thymocytes subsequently downregulated *Zbtb7b*, simultaneous with *Runx3* upregulation.

To confirm the co-expression of THPOK and RUNX3 in CD8-fated cells, we performed intracellular flow cytometry using OT-I TCRtg mice, an MHCI-specific transgenic model with a particularly prominent CD4+CD8low population (Lundberg et al., 1995). Wild-type and MHCII-specific OT-II transgenic model were included for comparison (Figure S4B-E). While THPOK and RUNX3 are generally expressed on distinct thymocyte populations, we observed a small population of co-expressing cells amongst positively selecting OT-I thymocytes and wild-type thymocytes, which was missing from OT-II transgenic thymocytes (Figure S4B). The level of RUNX3 expression on THPOK+ OT-I thymocytes was substantially lower than on mature CD8+ thymocytes, but was significantly above background, as determined by fluorescence minus one (FMO) controls and staining of THPOK+ OT-II thymocytes (Figure S4C-E). Moreover, the THPOK+RUNX3+ population in OT-I thymocytes was predominantly CD4+CD8low (Figure S4B), as predicted by CITE-seq data (Figure 3E). Co-expression of CD4- and CD8-defining transcription factors in MHCI-specific CD4+CD8low cells suggests that they represent cells that have recently failed the CD4 audition and are transitioning to undergo CD8-lineage specification. Altogether, the temporal and lineage specific expression pattern of transcription factors and TCR target genes shown here provide strong evidence for an initial CD4 auditioning phase for both MHCI- and MHCII-specific thymocytes and suggest a role for TCR signaling in late CD8 lineage specification.

The expression of the CD4 and CD8 coreceptors plays an important role not only in defining the lineage of mature T cells, but also in transmitting the TCR signals that are necessary for thymocyte development (Germain, 2002). The dynamic pattern of coreceptor protein expression fits well with published data, including an initial dip in expression of both coreceptors (“double dull” stage (Lucas and Germain, 1996)) followed by a rise in CD4 and a continued decrease in CD8 (Figure 3F). Later in the CD8 lineage, CD8 expression increased along with a decrease in CD4 expression, resulting in a late transient DP stage (DP3) as the cells moved towards the CD8 SP phenotype. Differential expression of coreceptor RNA and lineage defining transcription factors was in good agreement with their known regulatory relationships (Taniuchi, 2016), with a significant difference in *Cd8a* RNA expression starting at pseudotime point 6, corresponding to the rise in *Zbtb7b,* and differential expression of *Cd4* RNA starting at pseudotime point 9, corresponding to preferential expression of *Runx3* in the CD8 lineage (Figure 3G).

The dynamic pattern of coreceptor expression and its impact on TCR signaling is a key factor in defining CD4 versus CD8 lineage commitment (Shinzawa et al., 2022). In CD4-fated cells, coreceptor expression remains relatively high and is not correlated with expression of TCR target genes *Cd69* and *Egr1* (Figure 3A,F). Instead, the gradual decline in TCR signal after pseudotime point 3 is likely due to negative feedback, including induction of *Dusp2/5* (Fig S3B), dual phosphatases that inhibit ERK signaling (Kovanen et al., 2008; Tanzola & Kersh, 2006). In contrast, in CD8-fated cells, the more rapid decline in TCR signaling during the first wave coincided with the decline in CD8 expression, as predicted by the kinetic signaling model (Singer et al., 2008). Moreover, the second rise in TCR signal (pseudotimes 8-10) correlates well with the rise in CD8 protein expression. Thus the impact of increased CD8 in facilitating MHCI recognition, together with other factors that increase thymocyte sensitivity to TCR signals at later stages of positive selection (Saini et al., 2010; Au-Yeung et al., 2014, Lutes et al., 2021) can help to explain the second TCR signaling wave.

Taken together, the timing of these events encompassing coreceptors, master regulators, and TCR signaling responses support a sequential model of CD4 and CD8 T cell development, summarized in Figure S5A. In this model, all positively-selected DP thymocytes begin the process of lineage commitment by auditioning for the CD4 lineage, as GATA3 upregulation is followed by THPOK induction and a drop in CD8 expression in both MHCII- and MHCI-specific thymocytes. Relatively sustained TCR signals in MHCII-specific thymocytes during an initial wave of TCR signaling locks in the CD4 fate, likely due to higher GATA3 expression and the activation of the THPOK positive autoregulation loop (Muroi et al., 2008; Wang et al., 2008), and accompanied by full repression of the *Cd8* gene by THPOK. In contrast, the more transient first wave of TCR signaling in MHCI-specific thymocytes leads to less sustained expression of GATA3 and THPOK. As CD8-fated cells continue to mature, the window for CD4 commitment closes and the window for CD8 lineage specification opens, perhaps due to general maturation-associated changes including the decline in E protein (E2A and HEB) transcription factor activity (Jones-Mason et al., 2012). Indeed, we observed a steady downregulation of the E protein genes *Tcf3* (E2A) and *Tcf12* (HEB) as well as transient induction of the E protein inhibitors *Id2* and *Id3* during development (Figure S3B). During this window, *Runx3* expression rises as a second wave of TCR signaling occurs, corresponding to CD8 lineage specification. Our pseudotime analyses and this model provide a useful framework for further mechanistic dissection of the drivers of the CD4 versus CD8 lineage decision.

### Gene expression differences between CD4-fated and CD8-fated cells implicate putative drivers of lineage commitment

To better understand the process of lineage commitment, we systematically investigated how differences emerge between the CD4 and CD8 lineages by performing totalVI differential expression tests between lineage-restricted thymocytes within equivalent units of pseudotime (Methods). There were no substantial differences in either RNA or protein expression between CD4 versus CD8-fated thymocytes at the early DP stages, and differentially expressed features gradually accumulated throughout maturation (Figure 4A; Table S7). This analysis resulted in a set of 302 genes that had significantly higher expression in MHCII-specific thymocytes (“CD4-DE”) and 397 genes that had significantly higher expression in MHCI-specific thymocytes (“CD8-DE”) in at least one pseudotime unit (Figure S5B-C). In this test, 92 genes were included in both lists when they showed higher expression in one lineage followed by higher expression in the other lineage at a later pseudotime (Figure S5B-C). The genes in each set were clustered by their expression across all cells in the corresponding lineage (Figure 4B-C; Tables S8 and S9). Inspection of mean gene expression of each cluster over pseudotime revealed characteristic temporal patterns, reflecting variation in gene expression over the course of maturation within each lineage (Figure 4B-D). For example, CD4-DE cluster 5 and CD8-DE cluster 1 showed a late divergence of expression of master regulator transcription factors and genes related to effector functions in their respective lineages (e.g., *Zbtb7b* and *Cd40lg* in MHCII-specific cells, and *Runx3* and *Nkg7* in MHCI-specific cells). Of particular interest were three clusters that showed significant enrichment for TCR target genes (hypergeometric test, Benjamini-Hochberg (BH)-adjusted P < 0.05; Methods): CD4-DE clusters 4 and 7, and CD8-DE cluster 3. CD4-DE cluster 7 and CD8-DE cluster 3 contained many of the same target genes, including the classic TCR target genes *Cd69* and *Egr1* (Figure 4B-D) and showed an early expression peak that was more sustained in CD4-fated relative to the CD8-fated cells, and a second later peak, specifically in CD8-fated cells. TCR target genes in the CD4-DE clusters 4 displayed a similarly early, single peak which, again, was more sustained for CD4-fated cells. As a possible explanation for the late increase in TCR signaling in MHCI-specific thymocytes, we find that CD8-DE clusters 0 and 4 exhibited increased expression in the CD8 lineage just before the second rise in TCR signaling and contained genes implicated in modulating TCR sensitivity. For example, cluster 0 contained *Cd8a*, which is required for MHCI recognition, and *Themis*, which modulates TCR signal strength during positive selection (Choi et al., 2017). CD8-DE cluster 4 contained ion channel component genes *Kcna2* and *Tmie*, which have been proposed to play a role in enhancing TCR sensitivity in thymocytes with low self-reactivity (Lutes et al., 2021).

**Figure 4:**
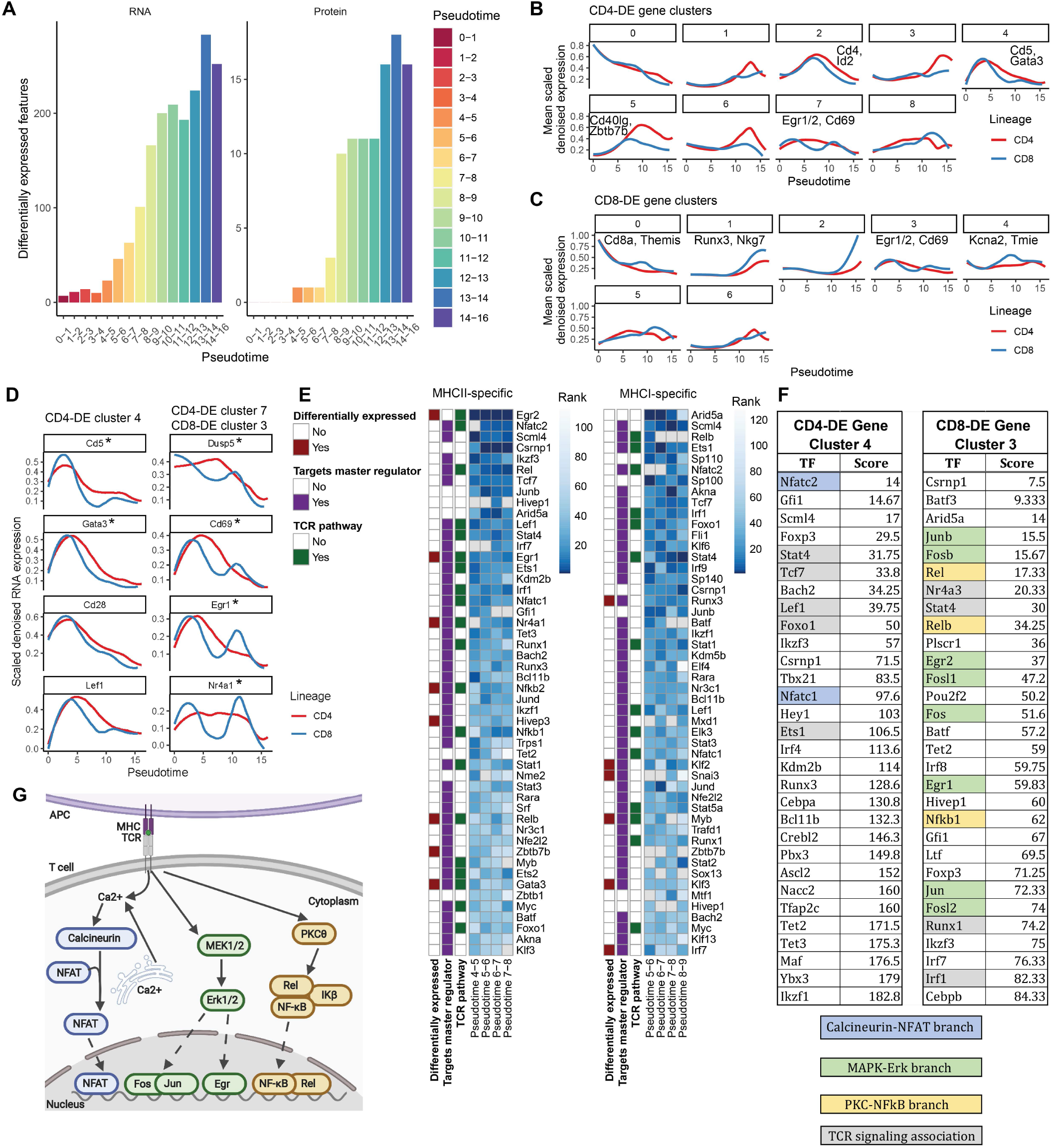
Gene expression differences between CD4-fated and CD8-fated cells implicate putative drivers of lineage commitment. **(A)** Number of differentially expressed features between the CD4 and CD8 lineages across pseudotime (Methods). **(B)** Genes (RNA) upregulated in the CD4 lineage relative to the CD8 lineage scaled per gene and clustered by the Leiden algorithm according to expression in the CD4 lineage. Expression over pseudotime per cluster is displayed as the mean of scaled totalVI denoised expression per gene for genes in a cluster, smoothed by loess curves. **(C)** Same as (B), but for genes upregulated in the CD8 lineage relative to the CD4 lineage, clustered according to CD8 lineage expression. Expression over pseudotime of selected TCR target genes. All genes are differentially expressed between the two lineages (with cluster membership indicated above) and are putative targets of *Nfatc2*, according to one or more of the ChEA3 analyses. Genes indicated with (*) are differentially expressed during the time windows used for ranking in (E). totalVI denoised expression values are scaled per gene and smoothed by loess curves. **(E)** Transcription factor enrichment analysis by ChEA3 for CD4-lineage-specific differentially expressed genes. Transcription factors are ranked by mean enrichment in the three pseudotime bins prior to *Zbtb7b* differential expression (between pseudotime 4-7; pseudotime 7-8 is for visualization and does not contribute to ranking). Gray indicates a gene detected in less than 5% of cells in the relevant population. “Differentially expressed” indicates significant upregulation in at least one of the relevant time bins. “Targets master regulator” indicates a transcription factor that targets either *Gata3*, *Runx3*, or *Zbtb7b* in ChEA3 databases. “TCR pathway” indicates membership in NetPath TCR Signaling Pathway, genes transcriptionally upregulated by TCR signaling, or genes with literature support for TCR pathway membership (Methods). Right panel shows the analyses for the CD8 lineage, with ranking by mean enrichment in the three pseudotime bins prior to *Runx3* differential expression (between pseudotime 5-8; pseudotime 8-9 is for visualization and does not contribute to ranking). **(F)** Transcription factor (TF) enrichment analysis for TCR target-enriched gene clusters CD4-DE clusters 4 and CD8-DE cluster 3 Figure 4B and C. The top 30 TFs enriched in the gene sets are shown. The full ChEA3 enrichment analysis is in Table S11 and S12. Colored boxes correspond to TFs activated by the respective branch of TCR signaling annotated in the inset diagram. Gray boxes indicate additional TFs associated with TCR signaling based on Netpath (Kandasamy et al., 2010), and as labeled in Figure 4E. **(G)** Schematic of the three major branches of the TCR signaling pathway: calcineurin-NFAT (blue), MEK-ERK (green), and PKC-NF-kB (orange).

To identify which transcription factors may influence lineage commitment, we focused on the pseudotime period just after differential gene expression was first detected and immediately upstream of master regulator expression (pseudotimes 4-7 for the CD4 lineage and pseudotimes 5-8 for the CD8 lineage). We performed transcription factor enrichment analysis with ChEA3 (Keenan et al., 2019), which identifies the transcription factors most likely to explain the expression of a set of target genes according to an integrated scoring across multiple information sources for potential regulatory activity, including ENCODE and ReMap ChIP-seq experiments (Dunham et al., 2012; Cheneby et al., 2020). We used differentially expressed genes between lineages in each unit of pseudotime as the target gene sets (Figure 4E; Tables S10 and S11; Methods) and ranked candidate transcription factors based on enrichment in each time unit and lineage. For each transcription factor, we also annotated whether there was a known association with TCR signaling (Kandasamy et al., 2010; Methods), evidence that the transcription factor regulates *Gata3*, *Zbtb7b*, or *Runx3* according to ChEA3 databases, and whether the transcription factor itself was differentially expressed at the relevant pseudotime stage. Focusing on MHCII-specific cells, several highly-ranked transcription factors were members of pathways associated with TCR signaling (e.g., *Egr2*, *Nfatc2*, *Egr1*, *Nfatc1*, and *Rel*). In particular, the top two ranked transcription factors in the CD4 lineage were *Egr2* and *Nfatc2*, which are known to lie downstream of two of the three main branches of the TCR signal transduction pathway: the extracellular signal-regulated kinase branch (hereafter called MEK-ERK) and calcineurin-NFAT branch, respectively (Figure 4G: Navarro and Cantrell, 2014; Hogquist and Jameson, 2014; Malissen et al., 2014; Chakraborty and Weiss, 2014). Notably, while the MEK-ERK branch plays a crucial role downstream of TCR signaling during positive selection (Sharp et al., 1997; Wilkinson and Kaye, 2001; Daniels et al., 2006; McNeil et al., 2005), and the NFKB pathway plays a role in late thymic development but is not required for positive selection (Xing et al., 2016), the role of the calcineurin-NFAT branch is less clear (Gallo et al., 2007).

To more closely explore how branches of the TCR signaling pathway are associated with divergent transcriptional regulation between the two lineages, we focused on genes in CD4-DE clusters 4 and 7 and CD8-DE cluster 3, which showed the greatest enrichment for TCR target genes (Figure 4B-D). Interestingly, while all three clusters showed an early peak that corresponded in pseudotime to the CD4 audition phase, CD4-DE cluster 4 exhibited more transient expression and lacked a prominent second peak in CD8-fated cells, implying that these genes might be regulated by a branch of the TCR signaling pathway that is selectively active early during the CD4 audition phase. This gene cluster contained *Gata3*, which plays a key role in CD4 fate by activating the CD4 master regulator *Zbtb7b* (Wang et al., 2008), and has been previously implicated as a target of the TCR-associated transcription factor NFAT (Gimferrer et al., 2011; Kandasamy et al., 2010; Scheinman and Avni, 2009). ChEA3 analysis of the genes in CD4-DE cluster 4 showed enrichment for NFAT family member *Nfatc2* (Figure 4F, Table S12), with *Gata3, Cd5, Id3, Cd28*, and *Lef1* all contributing to the enrichment score. In contrast, CD4-DE cluster 7 and CD8-DE cluster 3 showed enrichment for AP-1 transcription factors *Fosb* and *Junb*, NF-kB family members *Rel* and *Nfkb1/2*, and MEK-ERK target *Egr1* (Figures 4F and S5D; Table S13). This suggested that all three branches of the TCR signaling pathway participate during the CD4 audition phase, whereas the MEK-ERK and PKC-NF-kB branches, but not the calcineurin-NFAT branch, are active in the later specification of the CD8 lineage. Based on these data, together with the ranking of NFAT in driving early transcriptional differences between lineages (Figure 4E), as well as the dearth of information about the role of NFAT downstream of TCR signaling during positive selection, we chose to focus on the calcineurin-NFAT pathway for functional testing.

### Calcineurin-NFAT promotes commitment to the CD4 lineage via GATA3 induction

We used neonatal thymic slice cultures as an experimental system to study the impact of calcineurin inhibition on lineage commitment. Previous work has shown that genetic disruption of the calcineurin B1 regulatory subunit in thymocytes, or long term in vivo treatment with calcineurin inhibitors, leads to a developmental defect in DP thymocytes that obscures a possible role of calcineurin downstream of TCR signals during positive selection (Gallo et al., 2007). The neonatal thymic slice culture system circumvents this problem, because calcineurin can be inhibited after DP thymocytes are already present but prior to positive selection. Since mature SP thymocytes first appear shortly after birth in mice, we reasoned that neonatal thymic slice cultures would allow us to manipulate TCR signaling during a new wave of CD4 and CD8 SP development. We prepared thymic slices from postnatal day 1 mice and cultured slices for up to 96 hours on tissue culture inserts. We quantified populations of developing thymocytes based upon cell surface marker expression using flow cytometry (Figure S6A-B; Methods). As expected, at the start of the cultures we observed mostly DP thymocytes, whereas the frequencies of more mature populations including CD4+CD8low, CD4+ semimature (CD4 SM), CD4+ mature (CD4 SP), and CD8+ mature (CD8 SP) increased over time in culture (Figure S6B). Consistent with previously published results (Saini et al., 2010; Lucas et al., 1993) and our pseudotime analysis, CD8 T cell development was slightly delayed compared to that of CD4 T cells (Figure S6B). Thus, the neonatal slice system supports the development of both CD4 and CD8 lineage cells, providing an experimental time window to manipulate TCR signaling during CD4 and CD8 development with pharmacological inhibitors.

To directly test the involvement of calcineurin-NFAT during CD4 and CD8 thymic development, we inhibited calcineurin activity by adding Cyclosporin A (CsA) (Liu, 1993) to the neonatal slice cultures. Treatment of cultures for 96 hours at doses of CsA 200 ng/mL or below did not impact the relative size of most thymocyte populations while leading to a selective and dose dependent reduction in CD4+CD8low and CD4 SM thymocytes (Figure 5A). We observed a similar reduction in CD4+CD8low thymocytes in CsA-treated cultures of neonatal slices from MHCI-/- mice, while neonatal slice cultures from MHCII-/- mice had a reduced CD4+CD8low population that was not impacted by CsA (Figure S6C). Time course analyses of neonatal thymic slice cultures from wild-type mice showed that the reduction in CD4+CD8low and CD4 SM cells became significant after 72 and 48 hours of culture respectively (Figure 6B).

**Figure 5:**
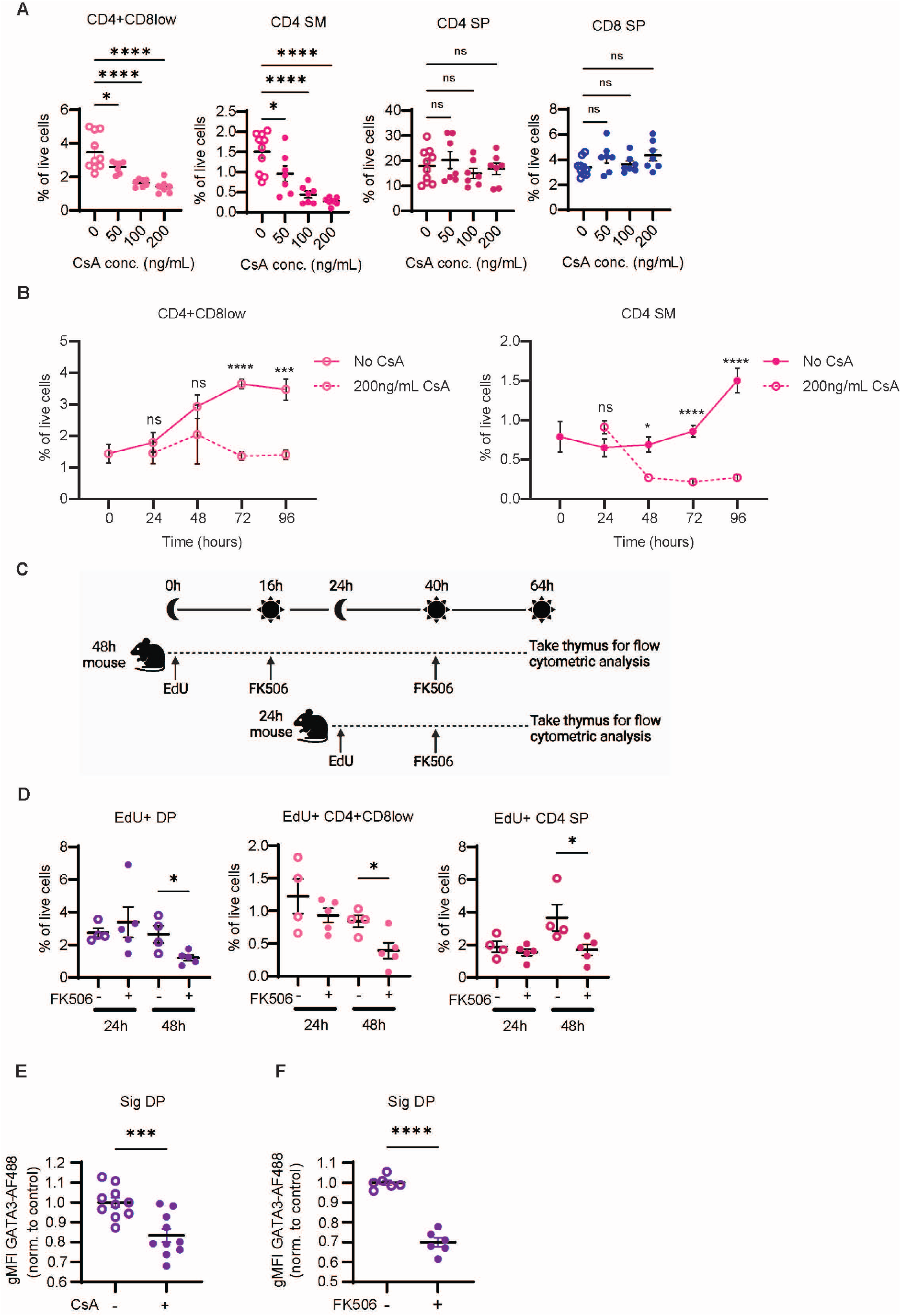
Inhibition of calcineurin blocks new CD4 SP development and GATA3 induction. Thymic tissue slices were prepared from postnatal (day 1) mice and cultured with the calcineurin inhibitor Cyclosporin A (CsA) at the indicated concentrations. Thymic slices were collected at indicated time points and analyzed via flow cytometry to quantify cell populations. Gating strategy is shown in Figure S6A. **(A)** Frequency (% of live cells) of CD4+CD8low, CD4 SM, CD4 SP, or CD8 SP in neonatal slices following culture with indicated concentrations of CsA for 96 hours. Data were compiled from 3 independent experiments with WT slices and analyzed using an ordinary one-way ANOVA. **(B)** Frequency (% of live cells) of CD4+CD8low and CD4 SM cells after 0, 24, 48, 72 and 96 hours of culture in medium alone or with 200 ng/mL CsA. Data were compiled from 9 independent experiments with WT slices and displayed as the mean ± standard error of the mean (SEM). For slices cultured with no CsA for 0 hours n=6, 24 hours n=9, 48 hours n=10, 72 hours n=22, 96 hours n=10. For slices cultured with 200 ng/mL CsA for 24 hours n=6, 48 hours n=7, 72 hours n=14, 96 hours n=7. Data was analyzed using an ordinary two-way ANOVA with multiple comparisons. **(C)** Schematic of in vivo EdU labeling and calcineurin blockade. AND TCRtg mice were injected with EdU to label proliferating thymocytes undergoing TCRβ selection and starting at 16 hours mice were treated with the calcineurin inhibitor FK506 daily for 24 or 48 hours. Thymocytes were analyzed by flow cytometry. **(D)** Frequency (% of live cells) of the indicated EdU+ thymocyte populations. Gating strategy is shown in Figure S6C. Each dot represents an individual mouse, and data are pooled from 3 independent experiments. **(E-F)** Geometric mean fluorescent intensity (gMFI) of GATA3 detected by intracellular flow cytometry staining in gated signaled (CD69+) DP thymocytes from neonatal thymic slice cultures after 48 hours of culture with no drug or with 200 ng/mL CsA **(E)** or after 72 hours with no drug or 6.3 ng/mL FK506 **(F)**. Data were compiled from 3 independent experiments with WT slices for (E) and 2 independent experiments with WT slices for (F). Each symbol on the graphs represents a thymic slice. Data was analyzed using an unpaired t test. NS is not significant, *p<0.05, **p<0.01, ***p<0.001, ****p<0.0001.

**Figure 6:**
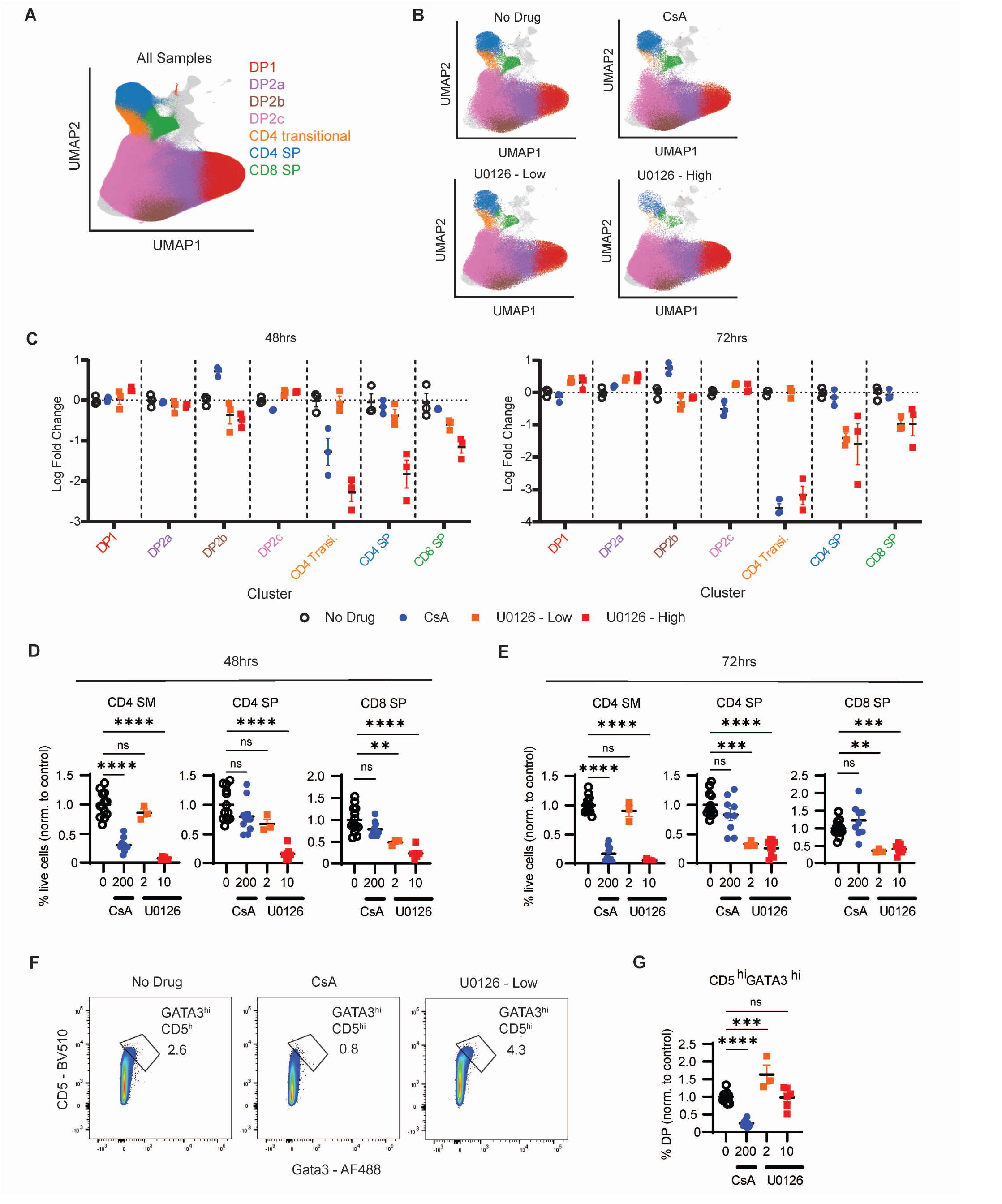
Calcineurin inhibition selectively impacts the CD4 audition. Thymic tissue slices were prepared from postnatal (day 1) mice and cultured with either calcineurin inhibitor CsA (200 ng/mL) or MEK inhibitor U0126 (2 μg/mL or 10 μg/mL). Thymic slices were collected at 48 or 72 hours, stained with fluorescent antibodies and analyzed by computational, multidimensional (A-C) and manual (D-G) gating. **(A)** UMAP plots with DP and SP populations highlighted. **(B)** UMAP plots for 48 hour time point separated by different drug treatment. **(C)** Scatter plots showing the log fold change in cell type proportions relative to no drug control for the indicated cell clusters for each condition. Each dot is a separate thymic slice, and data are normalized to the no drug condition. For (A–C), data is from one representative experiment out of two. **(D, E)** Manual gating using the gating strategy shown in Figure S6A. Plots show frequency (% of live cells), normalized to untreated control, of the indicated populations after 48 hours (D) or 72 hours of culture. **(F, G)** Upregulation of CD5 and GATA3 in gated DP thymocytes from 48 hour neonatal slice cultures. Representative flow cytometry plots **(F)** and compiled data **(G)** of gated DP thymocytes from the indicated conditions. For (D), (E), and (G), each dot is a thymic slice, and data are compiled from 3 independent experiments for CsA (200 ng/mL), one experiment for U0126 low (2 μg/mL), and 2 independent experiments for U0126 high (10 μg/mL).

The observation that low doses of CsA impacted the earliest subset of CD4 SP (CD4 SM), and the immediate precursor to CD4 SP (CD4+CD8low), but not the overall number of CD4 SP, suggested that many thymocytes in the neonatal culture may have already passed the stage requiring high calcineurin-NFAT signals prior to CsA addition. To test this hypothesis, we used in vivo EdU labeling of adult AND TCRtg mice to track cohorts of developing CD4 SP thymocytes during calcineurin blockade. For these experiments, we used the calcineurin inhibitor FK506, based on earlier literature showing that in vivo treatment with FK506 blocked positive selection without the loss of DP thymocytes observed with CsA treatment (Wang et al. 1995). AND TCRtg mice were injected with EdU to label proliferating thymocytes undergoing TCRβ selection and, starting 16 hours later, mice were injected daily with FK506 and analyzed 24 or 48 hours after the start of FK506 treatment (Figure 5C). In control mice that were injected with EdU but not FK506, the percentage of EdU-labeled CD4 SP increased from ∼2% to ∼4% between 24 and 48 hours, reflecting the conversion of labeled thymocytes from DP to CD4 SP during the time course (Figures 5D and S6D). Treatment of mice with FK506 had no significant impact on the overall percentage of DP or CD4 SP thymocytes (Figure S6E), and treatment for 24 hours had no significant impact on the number of EdU-labeled DP or CD4 SP (Figure 5D). However, 48 hours of in vivo exposure to FK506 led to a significant reduction in EdU-labeled CD4 SP, CD4+CD8low, and DP thymocytes (Figure 5D). Previous studies showed that long term loss of calcineurin by genetic ablation or lengthy (10 day) in vivo treatment with calcineurin inhibitors prevented normal ERK activation by DP thymocytes in response to TCR engagement (Gallo et al., 2007). To test whether this indirect impact of calcineurin blockade occurred in our short term (2 day) treatment, we performed TCR crosslinking and measured ERK activation by pERK staining of thymocytes from FK506-treated mice. We observed strong induction of pERK in DP thymocytes in response to TCR crosslinking in both 24- and 48-hour FK506-treated mice, similar to untreated controls (Figure S6F). Altogether, these data confirm that blocking calcineurin activation downstream of TCR signals during positive selection prevents new CD4 SP development, but does not interfere with already selected CD4 SP thymocytes.

The transcription factor *Gata3* is significantly upregulated in CD4-fated compared to CD8-fated thymocytes immediately prior to CD4 lineage commitment (Figure 3C-D), and was implicated by our ChEA3 analyses (Figure 4F,I) and prior studies (Gimferrer et al., 2011; Kandasamy et al., 2010; Scheinman and Avni, 2009) as a target of NFAT. Moreover, GATA3 induces the expression of the CD4 master regulator THPOK (Wang et al., 2008). Thus, we hypothesized that CsA treatment prevented new CD4 development by interfering with THPOK induction via GATA3. To test this hypothesis, we performed intracellular staining to quantify GATA3 expression in neonatal slice cultures treated with CsA for 48 hours (the time point at which we first observed a significant decrease in CD4 SM populations with CsA addition). In cultures with CsA, we observed a significant reduction in GATA3 expression in signaled (CD69+) DP thymocytes (Figure 5E).

To confirm these results, we also performed neonatal slice cultures in the presence of FK506. At moderate concentrations (0.4-6 ng/mL), the impact of FK506 was similar to CsA, causing a loss of CD4+CD8low and CD4 SM thymocytes without a significant loss of mature CD4 or CD8 SP (Figure S6G). Moderate concentrations of FK506 (6 ng/mL) in the neonatal system also led to a significant reduction in GATA3 protein expression in signaled (CD69+) DP (Figure 5F). Together these data validate predictions from our CITE-seq data that implicate the calcineurin/NFAT/GATA3 axis as a link between TCR signals downstream of MHCII recognition and commitment to the CD4 lineage.

### Calcineurin inhibition selectively impacts the CD4 audition

Transcription factor enrichment analyses of CITE-seq data suggested that different branches of the TCR signaling pathway exhibit distinct temporal and lineage-specific activity during positive selection. Specifically, a set of putative NFAT target genes were induced early during the CD4 audition, whereas putative targets of EGR and FOS/JUN transcription factors, which are regulated by the MEK-ERK pathway, peaked later in CD4-fated cells, and exhibited a second wave of induction in CD8-fated cells (Figure 4B-D, F-G). This suggested that the calcineurin-NFAT branch of the TCR signaling pathway might play a selective role in the CD4 audition, whereas the MEK-ERK branch may be required throughout CD4 and CD8 SP positive selection. To investigate this hypothesis, we compared the impact of calcineurin versus MEK inhibition in neonatal thymic slice cultures. To avoid off-target effects, we chose relatively low inhibitor concentrations based on titration experiments (Figure S7A). To define continuous developmental intermediates in an unbiased manner, we combined flow cytometric analyses of seven cell surface proteins and three lineage defining transcription factors, with a computational multidimensional gating approach by unsupervised clustering. These analyses identified populations containing mature CD4 and CD8 SP, CD4+CD8+ thymocytes, and a transitional CD4+CD8-population that largely overlapped with the CD4 semi-mature population identified by manual gating (colored areas in Figures 6A and S7B-C, and S8A-B). Smaller populations of αβTCR negative and mature, unconventional T cells were also identified (gray areas in Figure 6A-B, Figure S8A). Consistent with our results above (Figure 5), addition of 200 ng/mL of CsA for 48 or 72 hours led to significant reduction in the CD4 transitional population, with little impact on other populations (Figures 6B-C and S8C-D). In contrast, the MEK inhibitor U0126 at 10 μg/mL led to a loss of CD4 and CD8 SP as well as the CD4 transitional population, whereas a 5-fold lower concentration spared the transitional CD4 cells, while slightly reducing the CD4 SP and CD8 SP populations. Similar results were obtained upon manual gating of the data (Figure 6D-E). These data suggest that calcineurin inhibition impacts a relatively restricted temporal window during positive selection corresponding to the CD4 audition, whereas MEK inhibition impacts all stages of positive selection including CD8 specification.

In addition to the reduction in the CD4 transitional population, calcineurin inhibition by CsA impacted two of the DP populations (DP2b and DP2c) (Figure 6C). Interestingly, these populations were distinguished based on their relative levels of CD5 (Figure S7C), one of the putative NFAT targets based on transcription factor enrichment analyses (Figure 4E and Table S12). To further explore this difference, we used manual gating of DP thymocytes to compare the expression of CD5 and GATA3 (Figure 6F-G). We observed a reduction in the proportion of the CD5^high^GATA3^high^ population in DP thymocytes from 48-hour CsA treated samples, whereas this population was proportionally increased in the presence of low concentrations of U0126. These results are in line with our predictions from CITE-seq that NFAT activity targets the transient induction of a subset of TCR-target genes, including CD5 and GATA3. Taken together, these data support the conclusion that the calcineurin-NFAT branch of the TCR signaling pathway promotes strong induction of GATA3 during the CD4 audition, thus promoting CD4 fate commitment, whereas the MEK-ERK branch provides more general differentiation and survival signals throughout positive selection.

## Discussion

In this study, we applied single-cell multi-omic analysis to investigate the development of thymocytes into CD4 and CD8 T cells. By jointly analyzing the transcriptome and surface protein expression of thymocytes from both WT and lineage-restricted mice, we comprehensively defined the continuous changes over maturation in both lineages and linked them directly to intermediate populations traditionally defined by flow cytometry. We identified key lineage-specifying differences in gene and protein expression and determined the relative ordering of single cells along their differentiation trajectory by defining a pseudotime axis based on both the RNA and protein measurements. While previous studies indicated that phenotypically mature CD4 T cells appear earlier than CD8 T cells (Lucas et al., 1993; Saini et al., 2010; Karimi et al., 2021), our data show that this asynchrony corresponds to two distinct phases of differentiation: an initial CD4 “auditioning” phase in which both CD4 and CD8-fated cells undergo a parallel series of gene expression changes including up-regulation of the CD4-defining transcription factor *Zbtb7b* (encoding THPOK), followed by a CD8 lineage specification phase which corresponds to a 2^nd^ TCR signaling wave and overlaps with induction of the CD8-defining transcription factor *Runx3* (Figure S5A).

The availability of a high-resolution temporal map of gene and protein expression changes throughout positive selection and T cell lineage commitment provides a key resource for understanding mechanisms, as illustrated in the current study. We used this resource to pinpoint CD4-fated and CD8-fated thymocytes just prior to differential expression of the lineage-defining master transcription factors *Zbtb7b* (encoding THPOK) and *Runx3*, and to compare gene expression at this critical stage. In doing so, we identified the TCR/calcium/calcineurin-regulated transcription factor NFAT as a candidate to link TCR signaling upon MHCII recognition to upregulation of GATA3, THPOK, and CD4 lineage commitment. We also used this resource to define two distinct temporal waves of TCR signaling during positive selection: an early wave that coincides with the CD4 auditioning phase and is more sustained in MHCII-specific/CD4 lineage thymocytes compared to MHCI-specific/CD8 lineage thymocytes, and a later wave that overlaps with the CD8 specification stage and is specific for the CD8 lineage. While the inferred activity of some TCR-induced transcription factors, such as AP-1, EGR1/2, and NF-kB, occurred throughout positive selection, NFAT appeared to be most active during the CD4 audition phase, providing a potential link between TCR signals, *Gata3* induction, and the CD4 T cell fate. NFAT activity is required prior to positive selection to endow DP thymocytes with the ability to activate ERK upon TCR triggering (Gallo et al., 2007), but whether NFAT played a direct role downstream of TCR signaling during positive selection was unknown. Here we show that short term, low dose exposure to calcineurin inhibitors, conditions that did not impair ERK activation, prevented new development of mature CD4 thymocytes. As predicted by our CITE-seq analyses, we also observed that blockade of calcineurin led to decreased expression of GATA3 at the DP stage. These results are consistent with earlier studies implicating NFAT as a positive regulator of GATA3 (Gimferrer et al., 2011; Scheinman et al., 2009). Thus, computational analyses of CITE-seq data, as well as experimental manipulation, point to a role for NFAT downstream of TCR recognition of MHCII in driving CD4 lineage commitment.

The dynamic expression of coreceptors, TCR signaling targets, and transcription factors described here plays out over several days as thymocytes pass through windows of opportunity for CD4, then CD8, specification. Studies using synchronized systems for positive selection (Saini et al., 2010; Lutes et al., 2021) or BrdU-labeling of cohorts of thymocytes (Lucas et al., 1993) indicate that CD4 SP appear after 1-2 days of positive selection (corresponding to the CD4 audition: pseudotime points 0-6), whereas CD8 SP appear after 3 or more days (corresponding to CD8 lineage specification: pseudotime points 6-10). Thus, our data highlight the opportunity for multiple redundant mechanisms to guide the lineage choice. One proposed mechanism is a drop in CD8 expression leading to a more rapid decline in TCR signals that is proposed to signal thymocytes bearing MHCI-specific TCRs to adopt the CD8 fate, termed “kinetic signaling” (Singer et al., 2008). Consistent with this model, we observe a parallel decline in CD8 surface protein and TCR target gene expression in CD8-fated cells during the second half of the CD4 audition (pseudotimes 4-6). However, there are indications that this dip in CD8 expression is not an absolute requirement for CD8 fate specification. For example, while some MHCI-specific TCR transgenic models have a prominent CD4+CD8low thymocyte population (Lundberg et al., 1995), others do not (Lutes et al., 2021). In addition, mature CD8 T cells develop efficiently in the presence of constitutively expressed CD8 transgenes (see for example (Itano et al., 1996)). Thus, additional mechanisms contribute to directing MHCI-specific thymocytes to the CD8 fate.

Our data suggest that one such additional mechanism involves a late requirement for TCR signaling for CD8-fated cells. The observation of a second wave of TCR signaling coinciding with a rise in CD8 and *Runx3* expression in CD8-fated cells, suggests that late TCR signals may drive CD8 lineage specification. This notion fits with prior evidence for a prolonged requirement for TCR signaling for CD8 T cell development (Kisielow and Miazek, 1995; Liu and Bosselut, 2004; Au-Yeung et al., 2014; Sinclair and Seddon, 2014; McNeil et al., 2005), but are at odds with the kinetic signaling model, which invokes a complete loss of TCR signals and an exclusive role for cytokine signals during CD8 lineage specification (Singer et al., 2008). The second wave of TCR signaling may promote *Runx3* induction and survival of MHCI-specific thymocytes (Sinclair and Seddon, 2014). Such a mechanism likely contributes to the efficiency of lineage redirection in mice with reversed expression of coreceptor genes (Shinzawa et al., 2022), since expression of CD4 during the late “CD8 specification” phase would favor the development of redirected MHCII-specific CD4+ cytotoxic T cells.

Another proposed mechanism to help direct CD4/CD8 lineage commitment posits that stronger TCR signals during positive selection favor the CD4 fate (Karimi et al., 2021; Itano et al., 1996). While our data do not directly address this question, the notion of a limited developmental/temporal window for CD4 commitment, together with the greater ability of CD4 to recruit the tyrosine kinase LCK to the TCR complex, and indications that MHCII recognition generates inherently stronger signals than MHCI recognition (Moran et al., 2011), suggest that thymocytes bearing MHCII specific TCRs may accumulate TCR signals more rapidly during the CD4 audition, and thus are more likely to commit to the CD4 lineage. Data from coreceptor gene reversed mice (Shinzawa et al., 2022) are fully consistent with this possibility, since CD8+ MHCI-specific helper lineage T cells, which would be expected to experience weaker signals due to the use of the CD8 coreceptor, appear to express reduced levels of THPOK in the thymus and do not accumulate to normal numbers in the periphery.

Although CD8-fated cells undergo lineage specification late during positive selection, there are indications that they require TCR signals both early and late during their development (Kisielow and Miazek, 1995; Liu and Bosselut, 2004; Au-Yeung et al., 2014; Sinclair and Seddon, 2014). For example, inhibition of the TCR downstream kinase ZAP70 in any 12-hour window during a synchronous wave of positive selection prevented the emergence of mature CD8 SP thymocytes (Au-Yeung et al., 2014), implying that both the first and second waves of TCR signals are required for their development. Early TCR signals may promote survival or general maturation, thus allowing CD8-fated cells to progress to the stage where they become competent to undergo lineage specification. While it is clear that CD8 SP development requires TCR signals both early and late, whether this includes a specific requirement for calcineurin-NFAT remains unclear. While relatively low concentrations of calcineurin inhibitors blocked the development of early CD4 SP thymocytes without impairing CD8 SP development (Figures 5-6), we cannot exclude that a complete loss of calcineurin would block CD8SP. Future studies using synchronous models for positive selection combined with precisely timed genetic ablation approaches, such as targeted protein destabilization (Nabet et al., 2018) should help to address this question.

While in this study we focused our analysis on CD4/CD8 lineage commitment, it also serves to demonstrate the broader utility of using paired single-cell RNA and protein expression for studying developmental systems. The simultaneous measurement of RNA and protein not only allowed us to track the differences in relative timing of RNA and protein expression events, but also enabled more rigorous identification of cell populations isolated by FACS for further analysis, such as intracellular transcription factor staining or chromatin profiling. We also demonstrated that in silico gating of CITE-seq protein data can inspire gating strategies for fluorescence-based flow cytometry. The CITE-seq method is currently limited to the measurement of surface proteins (Stoeckius et al., 2017), but development of emerging methods such as inCITE-seq (Chung et al., 2021) to facilitate the simultaneous measurement of RNA, surface proteins, and large panels of intracellular proteins could greatly enhance the ability to generate hypotheses about molecular pathway activity, gene regulatory networks, and transcription and translation dynamics. Furthermore, since thymocytes actively traverse the thymic cortex and medulla over the course of their development, imaging could provide a valuable dimension to our current understanding of the thymocyte developmental timeline (Germain et al., 2012). Future work that integrates spatial genomic measurements with the transcriptomic and surface protein profiles generated in this study could inform how a cell’s micro-environment and physical motility might influence and reflect key aspects of the thymocyte developmental trajectory.

## Acknowledgments

We thank BioLegend Inc. and their proteogenomics team, especially Kristopher Nazor, Bertrand Yeung, Andre Fernandes, Qing Gao, Hong Zhang, and John Ma, for providing reagents and expertise and for help with sample preparation, library generation, and sequencing for a portion of the CITE-seq libraries used in this study as well as helpful discussions regarding analysis and totalVI. We thank the Cancer Research Lab Flow Cytometry Core Facilities at UC Berkeley, including Hector Nolla and Alma Valeros for their help operating cell sorters. We thank the UC Berkeley Functional Genomics Lab, especially Justin Choi. We thank Silvia Ariotti for insightful early discussions, and Adam Gayoso for helpful discussions on the application of totalVI. We would also like to thank Shiao Wei Chan and Kathya Arana for technical assistance. We thank Christina Usher for artwork. We thank members of the Streets, Yosef, and Robey laboratories for providing helpful feedback. Research reported in this manuscript was supported by the NIGMS of the National Institutes of Health under award number R35GM124916 (A.S), the NIAID of the National Institutes of Health under award number AI145816 (E.A.R., A.S., N.Y.), award number AI064227 (E.R.), and award number AI100829 (L.L.M.), the Chan Zuckerberg Foundation Network under grant number 2019-02452 (N.Y.) and the National Institutes of Mental Health under grant number U19MH114821 (N.Y.). Z.S. was supported by the National Science Foundation Graduate Research Fellowship and the Siebel Scholars award. N.Y. was supported by the Koret-Berkeley-Tel Aviv Initiative in Computational Biology. A.S. is a Pew Scholar in the Biomedical Sciences, supported by the Pew Charitable Trusts. A.S. and N.Y. are Chan Zuckerberg Biohub investigators.

## Author Contributions

Z.S. led the study with input from E.A.R., A.S., L.L.M., L.K.L., and N.Y. Z.S. and L.K.L. performed CITE-seq experiments. T.H. contributed towards sequencing and data processing of the cDNA and ADT CITE-seq libraries. D.A.A., L.L.M., J.Y.K., and I.B designed, performed, and analyzed thymic development and flow cytometry experiments with input from all authors. Z.S. designed and implemented analysis methods with input from all authors. D.A.A., C.E., L.L.M., J.Y.K., and I.B., and E.A.R. analyzed flow cytometry data with input from Z.S. Z.S., L.L.M., A.S., N.Y., and E.A.R wrote the manuscript. E.A.R., A.S., and N.Y. supervised the work.

## Methods

### CITE-seq on mouse thymocytes

#### Mice

All animal care and procedures were carried out in accordance with guidelines approved by the Institutional Animal Care and Use Committees at the University of California, Berkeley and at BioLegend, Inc. WT (B6) (C57BL/6, Stock No.: 000664), β2M-/- (B6.129P2-B2m^tm1Unc^/DcrJ, Stock No.: 002087; referred to as MHCI-/-), OT-I (C57BL/6-Tg(TcraTcrb)1100Mjb/J, Stock No.: 003831), and OT-II (B6.Cg-Tg(TcraTcrb)425Cbn/J, Stock No.: 004194) were obtained from The Jackson Laboratory. MHCII-/- (I-Aβ-/-) mice have been previously described (Grusby et al., 1991). AND TCRtg RAG1-/- mice and F5 TCRtg RAG1-/- mice were generated by crossing AND TCRtg (B10.Cg-Tg(TcrAND)53Hed/J, Jax Stock No.: 002761; (Kaye et al., 1989)) and F5 TCRtg (C57BL/6-Tg(CD2-TcraF5,CD2-TcrbF5)1Kio; (Mamalaki et al., 1992)) mice with RAG1-/- mice (Rag1-/-B6.129S7-Rag1tm1Mom) as previously described by (Au-Yeung et al., 2014)). All mice used in CITE-seq experiments were females between four and eight weeks of age. Samples are further described in Table S2. Mice were group housed with enrichment and segregated by sex in standard cages on ventilated racks at an ambient temperature of 26 °C and 40% humidity. Mice were kept in a dark/light cycle of 12 h on and 12 h off and given access to food and water ad libitum.

#### Cell preparation

Mice were sacrificed, and thymi were harvested, placed in RPMI + 10% FBS medium on ice, mechanically dissociated with a syringe plunger, and passed through a 70 μm strainer to generate a single-cell suspension.

#### Antibody panel preparation

We prepared a panel containing 111 antibodies (TotalSeq-A mouse antibody panel 1, BioLegend, 900003217), which are enumerated in Table S1. Immediately prior to cell staining, we centrifuged the antibody panel for 10 minutes at 14,000 g to remove antibody aggregates. We then performed a buffer exchange on the supernatant using a 50 kDa Amicon spin column (Millipore, UFC505096) following the manufacturer’s protocol to transfer antibodies into RPMI + 10% FBS.

#### Cell sorting

To enrich for positively-selecting thymocytes in MHC-deficient and some WT samples (Table S2), live, single, TCRβ+CD5+ thymocytes were sorted by FACS. We took advantage of the fact that cells were already stained with TotalSeq (oligonucleotide-conjugated) antibodies and therefore designed oligonucleotide-fluorophore conjugates complementary to the TotalSeq barcodes (5’-CACTGAGCTGTGGAA-AlexaFluor488-3’ for CD5; 5’-TCCCATAGGATGGAA-AlexaFluor647-3’ for TCRb). Prior to cell staining, the TotalSeq antibody panel was mixed with oligonucleotide-fluorophore conjugates in a 1:1.5 molar ratio. This mixture was incubated for 15 minutes at room temperature to allow for oligonucleotide hybridization, and then transferred to ice. Cells were then stained with the antibody/oligonucleotide-fluorophore mixture according to the TotalSeq protocol. Cells were stained, washed, and resuspended in RPMI + 10% FBS to maintain viability. Cells were sorted using a BD FACSAria Fusion (BD Biosciences).

#### CITE-seq protocol and library preparation

The CITE-seq experiment was performed following the TotalSeq protocol. Cells were stained, washed, and resuspended in RPMI + 10% FBS to maintain viability. We followed the 10X Genomics Chromium Single Cell 3′ v3 protocol to prepare RNA and antibody-derived-tag (ADT) libraries (Zheng et al., 2017).

#### Sequencing and data processing

RNA and ADT libraries were sequenced with either an Illumina NovaSeq S1 or an Illumina NovaSeq S4. Reads were processed with Cell Ranger v.3.1.0 with feature barcoding, where RNA reads were mapped to the mouse mm10–2.1.0 reference (10X Genomics, STAR aligner (Dobin et al., 2013)) and antibody reads were mapped to known barcodes (Table S1). No read depth normalization was applied when aggregating samples.

### CITE-seq data preprocessing

Prior to analysis with totalVI, we performed preliminary quality control and feature selection on the CITE-seq data. Cells with a high percentage of UMIs from mitochondrial genes (> 15% of a cell’s total UMI count) were removed. We also removed cells expressing < 200 genes, and retained cells with protein library size between 1,000 and 10,000 UMI counts. We removed cells in which fewer than 70 proteins were detected of the 111 measured in the panel. An initial gene filter removed genes expressed in fewer than four cells. The top 5,000 highly variable genes (HVGs) were selected by the Seurat v3 method (Stuart et al., 2019) as implemented by scVI (Lopez et al., 2018). In addition to HVGs, we also selected genes encoding proteins in the measured antibody panel and a manually selected set of genes of interest. After all filtering, the CITE-seq dataset contained a total of 72,042 cells, 5,125 genes, and 111 proteins.

### CITE-seq data analysis with totalVI

#### totalVI modeling of all CITE-seq data

We ran totalVI on CITE-seq data after filtering (described above), using a 20-dimensional latent space, a learning rate of 0.004, and early stopping with default parameters. Each 10X lane was treated as a batch. When generating denoised gene and protein values, we applied the *transform_batch* parameter (Gayoso et al., 2021) to view all denoised values in the context of WT samples.

#### Cell annotation

We stratified cells of the thymus into cell types and states based on the totalVI latent space, taking advantage of both RNA and protein information. We first clustered cells in the totalVI latent space with the Scanpy (Wolf et al., 2018) implementation of the Leiden algorithm (Traag et al., 2019) at resolution 0.6, resulting in 18 clusters. We repeated this approach to subcluster cells. We used Vision (DeTomaso et al., 2019) with default parameters for data exploration. Subclusters were manually annotated based on curated lists of cell type markers (Gayoso et al., 2021; Hogquist et al., 2015), resulting in 20 annotated clusters (excluding one cluster annotated as doublets). We visualized the totalVI latent space in two dimensions using the Scanpy (Wolf et al., 2018) implementation of the UMAP algorithm (Becht et al., 2019).

#### Differential expression testing of annotated cell types

We conducted a one-vs-all differential expression test between all annotated cell types, excluding clusters annotated as doublets or dying cells. We identified cell type markers by filtering for significance (log(Bayes factor) > 2.0 for genes, log(Bayes factor) > 1.0 for proteins), effect size (median log fold change (LFC) > 0.2 for both genes and proteins), and the proportion of expressing cells (detected expression in > 10% of the relevant population for genes), and sorting by the median LFC. For marker visualization, we selected the top four (if existing) differentially expressed genes and proteins per cell type, arranged by the cell type in which the LFC was highest.

#### totalVI modeling of positively-selecting thymocytes

To further analyze thymocyte populations with a focus on positively-selected cells, we selected the following annotated clusters: Signaled DP, Immature CD4, Immature CD8, Mature CD4, Mature CD8, Interferon signature cells, Negative selection (wave 2), and Treg. With an interest in the variation within thymocyte populations (rather than all cells in the thymus), we selected the top 5,000 HVGs in this subset, as well as genes encoding proteins in the measured antibody panel and a manually selected set of genes of interest. This resulted in a CITE-seq dataset containing 35,943 cells, 5,108 genes, and 111 proteins. We ran totalVI on this subset dataset and generated denoised values as described above. We performed Leiden clustering and visualized the totalVI latent space in two dimensions using UMAP as described above.

#### Cell filtering of positively-selecting thymocytes on the CD4/CD8 developmental trajectory

After visualizing the totalVI latent space of the thymocyte subset, we applied additional filters to restrict to cells on the CD4/CD8 developmental trajectory. We used two resolutions of Leiden clustering (0.6 and 1.4) and sub-clustering as described above to identify and remove clusters of negatively selected cells, Tregs, gamma-delta-like cells, mature cycling cells, and outlier clusters of doublets, interferon signature cells, and CD8-transgenic-specific outlier cells. After filtering, this dataset contained 29,408 cells that were used for downstream analysis. Differential expression testing of positively-selecting thymocytes using pseudotime information is described below.

### Pseudotime inference

#### Pseudotime inference with Slingshot

Slingshot (Street et al., 2018) was selected for pseudotime inference based on its superior performance in a comprehensive benchmarking study (Saelens et al., 2019). Slingshot pseudotime was derived from the UMAP projection of the totalVI latent space. The starting point was assigned to DP cells, and two endpoints were assigned to mature CD4 and CD8 T cells. Slingshot pseudotime derived from the full 20-dimensional totalVI latent space was highly correlated with that from the 2-dimensional space (Figure S2A), supporting our use of the 2D-derived pseudotime values for ease of visualization and analysis.

#### Lineage assignment

Initial lineage assignment of cells was made on the basis of their genotype (CD4 lineage for MHCI-/-, AND, and OT-II mice, CD8 lineage for MHCII-/-, F5, and OT-I mice, unassigned for WT mice). However, small numbers of cells in MHC-deficient and TCR transgenic mice develop along the alternative lineage (particularly in TCR transgenics that are *Rag* sufficient, which might express an endogenous TCR in addition to the transgenic TCR). We therefore added an additional filter of Slingshot lineage assignment weight > 0.5. Cells with a Slingshot lineage assignment weight of < 0.5 along the expected lineage based on genotype were excluded from the remaining pseudotime-based analysis.

### In silico flow cytometry and gating

To perform in silico flow cytometry, totalVI denoised protein counts were log-transformed and visualized in biaxial-style scatter plots. Gates in biaxial plots were determined based on contours of cell density. An approximate alignment of gated populations to pseudotime was generated by identifying thresholds classifying adjacent populations in pseudotime by maximizing the Youden criteria.

### Adult thymocyte population analysis with fluorescence-based flow cytometry

#### Mice

All experiments were approved by the University of California, Berkeley Animal Use and Care Committee. All mice were bred and maintained under pathogen-free conditions in an American Association of Laboratory Animal Care-approved facility at the University of California, Berkeley. WT (C57BL/6, Stock No.: 000664) and β2M-/- (B6.129P2-B2m^tm1Unc^/DcrJ, Stock No.: 002087) were obtained from The Jackson Laboratory. MHCII-/- (I-Aβ-/-) mice have been previously described (Grusby et al., 1991). For thymocyte population analysis in adult mice, six to eight week-old animals were used. Thymi were analyzed from eight mice per genotype (four male and four female).

#### Flow cytometry

Thymi were mechanically dissociated into a single-cell suspension, depleted of red blood cells using ACK Lysis Buffer (0.15M NH_4_Cl, 1mM KHC_3_, 0.1mM Na_2_EDTA). Cells were filtered, washed, and counted before being stained with a live/dead stain; Zombie NIR Fixable Viability Kit (Biolegend). Samples were blocked with anti-CD16/32 (2.4G2) and stained with surface antibodies against CD4, CD8, TCRβ, CD5, CD69, and CD127 (IL-7Ra) in FACS buffer (1% BSA in PBS) containing Brilliant Stain Buffer Plus (BD Biosciences). Intracellular staining for GATA3, THPOK, and RUNX3 was performed using the eBioscience FOXP3/Transcription Factor Staining Buffer Set (Thermo Fisher). All antibodies were purchased from BD Biosciences, Biolegend, or eBiosciences. Single-stain samples and fluorescence minus one (FMO) controls were used to establish PMT voltages, gating and compensation parameters. Cells were processed using a BD LSRFortessa or BD LSRFortessa X20 flow cytometer and analyzed using FlowJo software (Tree Star). Gates defining all populations were based on in silico-derived gates for all described proteins with the exception of CD127 in the CD4+ SM, CD8+ SM, CD4+ Mat, and CD8+ Mat populations. In these cases, the CD127 fluorescent antibody did not have comparable sensitivity to the CD127 CITE-seq measurement and was therefore excluded.

### Differential expression analysis of positively-selecting thymocytes with totalVI

#### Testing for temporal features

Temporal features (i.e., features that are differentially expressed over time) were determined by a totalVI one-vs-all DE test within each lineage between binned units of pseudotime. DE criteria (as above) included filters for significance (log(Bayes factor) > 2.0 for genes, log(Bayes factor) > 1.0 for proteins), effect size (median log fold change > 0.2 for both genes and proteins), and the proportion of expressing cells (detected expression in > 5% of the relevant population for genes). Top temporal genes were selected as the unique set among the top three differentially expressed genes per time that were differentially expressed in both lineages.

#### Testing for differences between lineages

Differences between lineages were determined by a totalVI within-cluster DE test, where clusters were binned units in pseudotime and the condition was lineage assignment (i.e., cells within a given unit of pseudotime were compared between lineages). Criteria for DE were the same as above.

#### Clustering of differentially expressed genes

To cluster differentially expressed genes into patterns, totalVI denoised gene expression values were standard scaled, reduced dimensions across cells using PCA, and clustered genes using the Leiden algorithm (Traag et al., 2019) as implemented by Scanpy (Wolf et al., 2018). For features differentially expressed between lineages, the genes upregulated within a lineage were clustered according to expression within the lineage in which they were upregulated.

#### Enrichment of TCR signaling in gene clusters

A hypergeometric test (phyper) was performed to test for enrichment of TCR signaling in differentially-expressed gene clusters. TCR signaling genes were compiled from Netpath (Kandasamy et al., 2010) and a set of genes activated upon stimulation in DP thymocytes (Mingueneau et al., 2012). The background set included all genes considered in DE analysis. P-values were adjusted by the Benjamini-Hochberg procedure.

### Transcription factor enrichment analysis

#### ChEA3 analysis

To perform transcription factor enrichment analysis with ChEA3 (Keenan et al., 2019), we first selected target gene sets as genes differentially upregulated in one lineage relative to the other in each unit of pseudotime, filtered for significance (log(Bayes factor) > 2.0), effect size (median log fold change > 0.2), and detected expression in > 5% of the population of interest. For each target gene set, transcription factors were scored for enrichment by the integrated mean ranking across all ChEA3 gene set libraries (MeanRank) based on the top performance of this ranking method (Keenan et al., 2019). ChEA3 analysis on gene clusters was performed as above, but using gene clusters as the target gene set.

#### Ranking of candidate transcription factors

To generate an overall ranking of transcription factors for their likely involvement in CD4/CD8 lineage commitment, we focused on enrichment in the three units of pseudotime prior to master regulator differential expression in each lineage (i.e., in the CD4 lineage, the relevant pseudotime units are 4, 5, and 6, prior to the differential expression of *Zbtb7b* differential expression at pseudotime 7; in the CD8 lineage, the relevant pseudotime units are 5, 6, and 7, prior to the differential expression of *Runx3* at pseudotime 8). We excluded the pseudotime unit containing master regulator differential expression from the ranking, as genes differentially expressed at this time could be the result of the master regulator itself enforcing lineage-specific changes rather than the factors driving initial commitment to a lineage. The pseudotime unit containing master regulator differential expression is included in Figure 4E for visualization, but did not contribute to the ranked order of transcription factors. We also excluded earlier units of pseudotime since these times included very few (< 15) significantly different genes between the lineages. Finally, pseudotime bins in which a transcription factor was not expressed in at least 5% of the population of interest, did not contribute towards that transcription factor’s ranking. The overall ranking of candidate driver transcription factors was then generated by taking the mean of ranks across the relevant pseudotime units.

#### TCR signaling pathway involvement

Transcription factors were annotated by whether they had a known association with TCR signaling. A list of molecules involved in TCR signaling were curated from the NetPath database of molecules involved in the TCR signaling pathway and the NetPath database of genes transcriptionally upregulated by the TCR signaling pathway (Kandasamy et al., 2010). Additional genes related to TCR signaling were curated from literature sources (Shao et al., 1997; Wong et al., 2014; Lopez-Rodriguez et al., 2015; Hedrick et al., 2013; Wang et al., 2010). Transcription factors were also annotated by whether they were known to target either *Gata3*, *Zbtb7b*, or *Runx3* according to ChEA3 databases (i.e., *Gata3*, *Zbtb7b*, or *Runx3* appeared in the Overlapping Gene list for the transcription factor of interest in any ChEA3 query).

### Neonatal thymic slice experiments

#### Mice

All experiments were approved by the University of California, Berkeley Animal Use and Care Committee. All mice were bred and maintained under pathogen-free conditions in an American Association of Laboratory Animal Care-approved facility at the University of California, Berkeley. WT (C57BL/6, Stock No.: 000664). For neonatal thymic slice experiments, postnatal day 1 (P1) mice were used.

#### Thymic slices

We prepared thymic slices from postnatal day 1 mice, a time point that allows us to track a synchronous wave of developing CD4 and CD8 thymocytes since T cells in mice do not develop until birth. Thymic slices were prepared as previously described (Dzhagalov et al., 2012; Ross et al., 2015), with minor modifications to adjust for the smaller size of neonatal thymi compared to those of adults. Thymic lobes were dissected, removed of connective tissue, embedded in 4% low melting point agarose (GTG-NuSieve Agarose, Lonza) and sectioned into 500 μM slices using a vibratome (VT1000S, Leica). Slices were overlaid onto 0.4 μM transwell inserts (Corning, Cat. No.: 353090) and placed in a 6-well tissue culture plate with 1 mL of complete RPMI medium (RPMI-1640 (Corning), 10% FBS (Thermo), 100 U/mL penicillin/streptomycin (Gibco), 1X L-glutamine (Gibco), 55 µM 2-mercaptoethanol (Gibco). Slices were cultured for indicated periods of time at 37 °C, 5% CO_2_, before being prepared and analyzed by flow cytometry. For neonatal slice cultures containing Cyclosporin A (CsA; Millipore-Sigma, Cat. No.:239835), CsA was serially diluted to indicated concentrations (50-800 ng/mL) and added directly to the culture medium. FK506 (Tacrolimus; InvivoGen, Cat. No.: inh-fk5-5) and U0126 (InvivoGen, Cat. No.: tlrl-u0126) were serially diluted in indicated concentrations (0.39-6.3 ng/mL and 0.63-10 µg/mL, respectively) and added directly to culture medium.

#### Flow cytometry

Thymic slices were mechanically dissociated into a single-cell suspension, then filtered, washed and counted before being stained with a live dead/stain; Propidium Iodine (Biolegend), Ghost Violet 510 (Tonbo), Zombie NIR, or Zombie UV Fixable Viability Kit (Biolegend). Samples were blocked with anti-CD16/32 (2.4G2) and stained with surface antibodies against CD4, CD8, TCRβ, and CD69 in FACS buffer (1% BSA in PBS) containing Brilliant Stain Buffer Plus (BD Biosciences). Intracellular staining for GATA3, RUNX3, and THPOK was performed using the eBioscience FoxP3/Transcription Factor Staining Buffer Set (Thermo Fisher). All antibodies were purchased from BD Biosciences, Biolegend, or eBiosciences. Single-stain samples and fluorescence minus one (FMO) controls were used to establish PMT voltages, gating and compensation parameters. Cells were processed using a BD LSRFortessa or BD LSRFortessa X20 flow cytometer and analyzed using FlowJo software (Tree Star). Note that the DP3 population is difficult to detect in neonatal compared to adult thymocytes samples, therefore we did not include it in our gating strategy.

#### Computational multi-dimensional analyses of flow cytometry data

FCS files were loaded into python using flowIO. Compensation was performed using manually determined compensation values. Data was loaded into Scanpy (Wolf et al., 2018) for further processing. Permissive manual gating in python was performed using physical dimension (FSC-W, FSC-A, SSC-A) on manual inspection and dead cells were filtered out based on live/dead staining. Fluorescent channels were normalized to a range of [0, 1]. Clustering was performed using PARC (Stassen et al., 2020) with a *resolution_parameter= 1.5*, *keep_all_local_dist=False* and *jac_std_global=0.15*. This yielded 34 clusters. Clusters were merged based on manual inspections of all fluorescent channels and merging was not performed if at least one fluorescent channel was differentially expressed between two clusters. We used PAGA (Wolf et al., 2019) initialization for UMAP embedding. PAGA was computed using 30 nearest neighbors in expression space using cosine distance. For UMAP embedding, we used the following parameters: *n_neighbors=30, metric=’euclidean’, min_dist=0.3, init_pos=’paga’*. For display of proportional changes in cluster frequency we divided the number of cells in each cluster by the total number of cells in the respective sample. We divided those values by the mean over the proportion in the respective cluster in the no drug sample and took the logarithm of this ratio to yield the log fold enrichment of the respective cluster. Seaborn was used for visualization. All computational gates were validated by manual inspection in FlowJo.

### In vivo EdU labeling and calcineurin blockade in adult mice

AND TCRtg RAG1-/- mice (described above) were intraperitoneally injected with 2 mg EdU (ThermoFisher Scientific Cat. #A10044) in the evening. The next morning (16h later), mice were injected with 5 μg FK506 (Invitrogen Cat. #INH-FK5-5). Thymi were taken for flow cytometry 24 or 48h after FK506 was administered. Thymi were dissociated and 2 × 10^6 cells were surface stained for flow cytometry as described above. After surface staining, cells were split, and 1 × 10^6 were processed using Click-iT EdU Pacific Blue Flow Cytometry Assay Kit (Thermo Fisher Scientific Cat. #C10418). The other 1 × 10^6 were subjected to anti-CD3 crosslinking and pERK staining as described below. Flow cytometry and data analysis were performed as described above.

### In vitro TCR activation and staining for phosphorylated ERK (pERK)

1×10^6 surface stained thymocytes were washed and resuspended in approximately 240 μL of serum-free media. Per sample, 10 μL of anti-CD3e antibody (clone 145-2C11, Invitrogen #14-0031-85) was added to reach a final concentration of 20 mg/mL. Working quickly, 7 μL of anti-Armenian hamster IgG crosslinker (Jackson ImmunoResearch Laboratories #127-0051-160) was added to each sample, briefly vortexed, and placed in a 37°C water bath for 3 minutes for in vitro TCR activation. For fixation, 1:1 volume of 4% paraformaldehyde was added to each tube and incubated at room temperature for 10 minutes before being washed in PBS. Cells were resuspended in 900 μL of ice-cold methanol by gentle pipetting, and incubated on ice for 30 minutes. After three washes in PBS, cells were incubated at 4°C overnight pERK antibody (1:20 dilution, Biolegend Cat. #675504). Samples were washed and resuspended for analysis. Flow cytometry and data analysis were performed as described above.

#### Statistical analysis

Data were analyzed using Prism software (GraphPad). Comparisons were performed using an unpaired T test, one- or two-way analysis of variance, where indicated in the figure legends. For all statistical models and tests described above, the significance is displayed as follows; ns is not significant, *p<0.05, **p<0.01, ***p<0.001, ****p<0.0001.

#### Figures#

Figures were created using Adobe Illustrator software. Illustrations in Figures 4G were created using Biorender.com.

## Supplemental Tables

**Table S1: Antibodies used in this study.**

**Table S2: CITE-seq sample information.**

**Table S3: DE test results for totalVI one-versus-all DE test between annotated thymus populations.**

**Table S4: Lineage information by genotype.**

**Table S5: DE test results for totalVI DE test across pseudotime within the CD4 lineage.**

**Table S6: DE test results for totalVI DE test across pseudotime within the CD8 lineage.**

**Table S7: DE test results for totalVI DE test within pseudotime and between CD4 and CD8 lineages.**

**Table S8: Cluster assignments for genes upregulated in the CD4 lineage from the totalVI DE test within pseudotime and between CD4 and CD8 lineages.**

**Table S9: Cluster assignments for genes upregulated in the CD8 lineage from the totalVI DE test within pseudotime and between CD4 and CD8 lineages.**

**Table S10: ChEA3 results for the CD4 lineage by pseudotime.**

**Table S11: ChEA3 test results for the CD8 lineage by pseudotime.**

**Table S12: ChEA3 test results for the CD4 lineage by gene cluster.**

**Table S13: ChEA3 test results for the CD8 lineage by gene cluster.**

## Data and Code Availability

CITE-seq data have been uploaded to GEO. An accession number will be provided prior to publication. Code will be made available prior to publication.

## Supplemental Figures Titles and Legends

**Figure S1:**
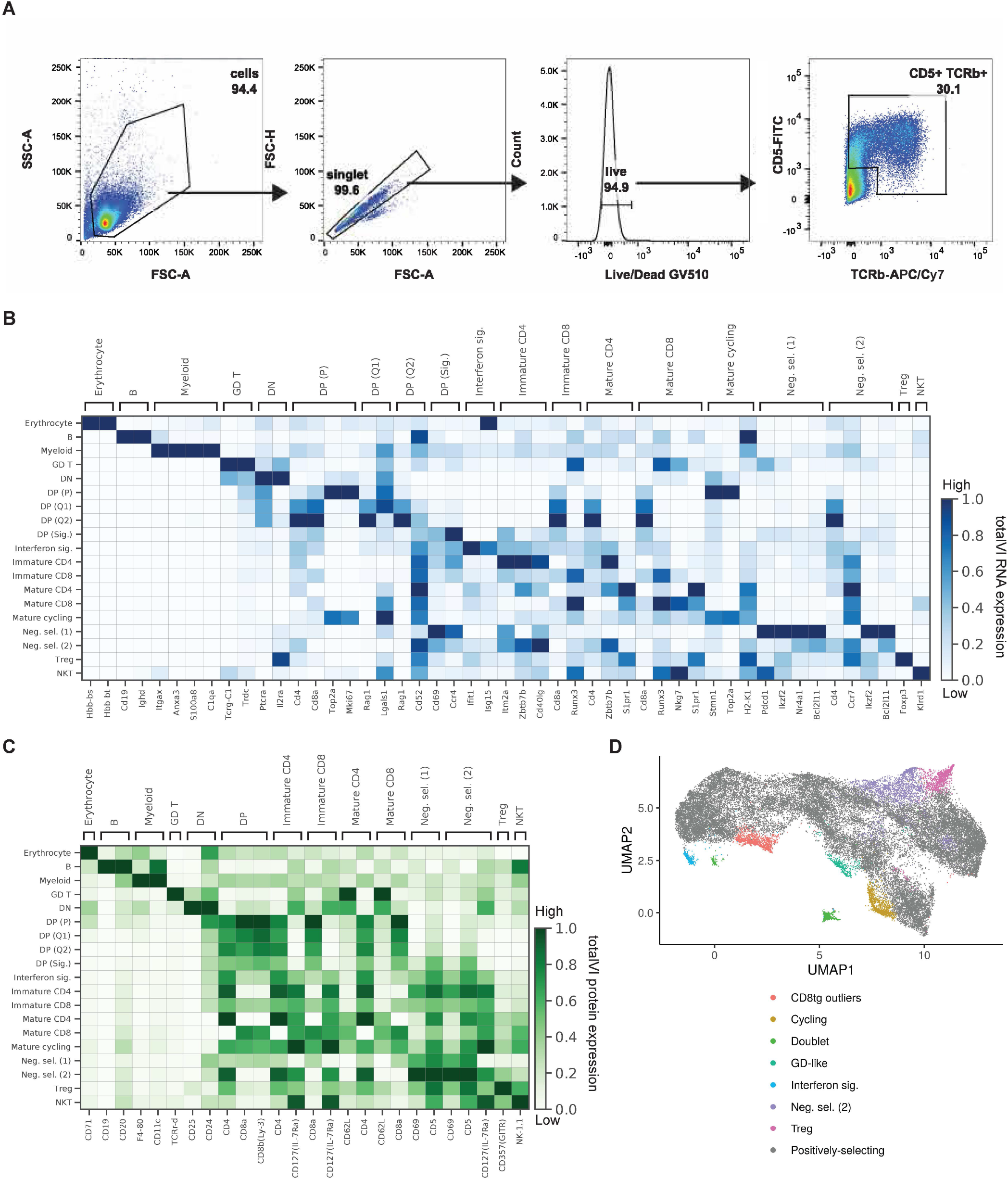
Sorting and characterization of positively-selecting thymocytes. **(A)** Representative FACS plots displaying gating strategy to sort thymocytes for CITE-seq. Cell populations were gated and sorted to include lymphocytes, exclude forward scatter doublets, include Ghost Dye Violet 510 Live/Dead stain negative (live cells), then on TCRβ+CD5+ to enrich for cells that were positively-selecting. **(B-C)** Heatmaps of manually selected cell type markers for **(B)** RNA and **(C)** proteins. Values are totalVI denoised expression. **(D)** UMAP plot of totalVI latent space from positively-selected thymocytes before filtering indicating annotated populations that were retained (positively-selecting thymocytes) or removed (all other populations) from downstream analysis.

**Figure S2:**
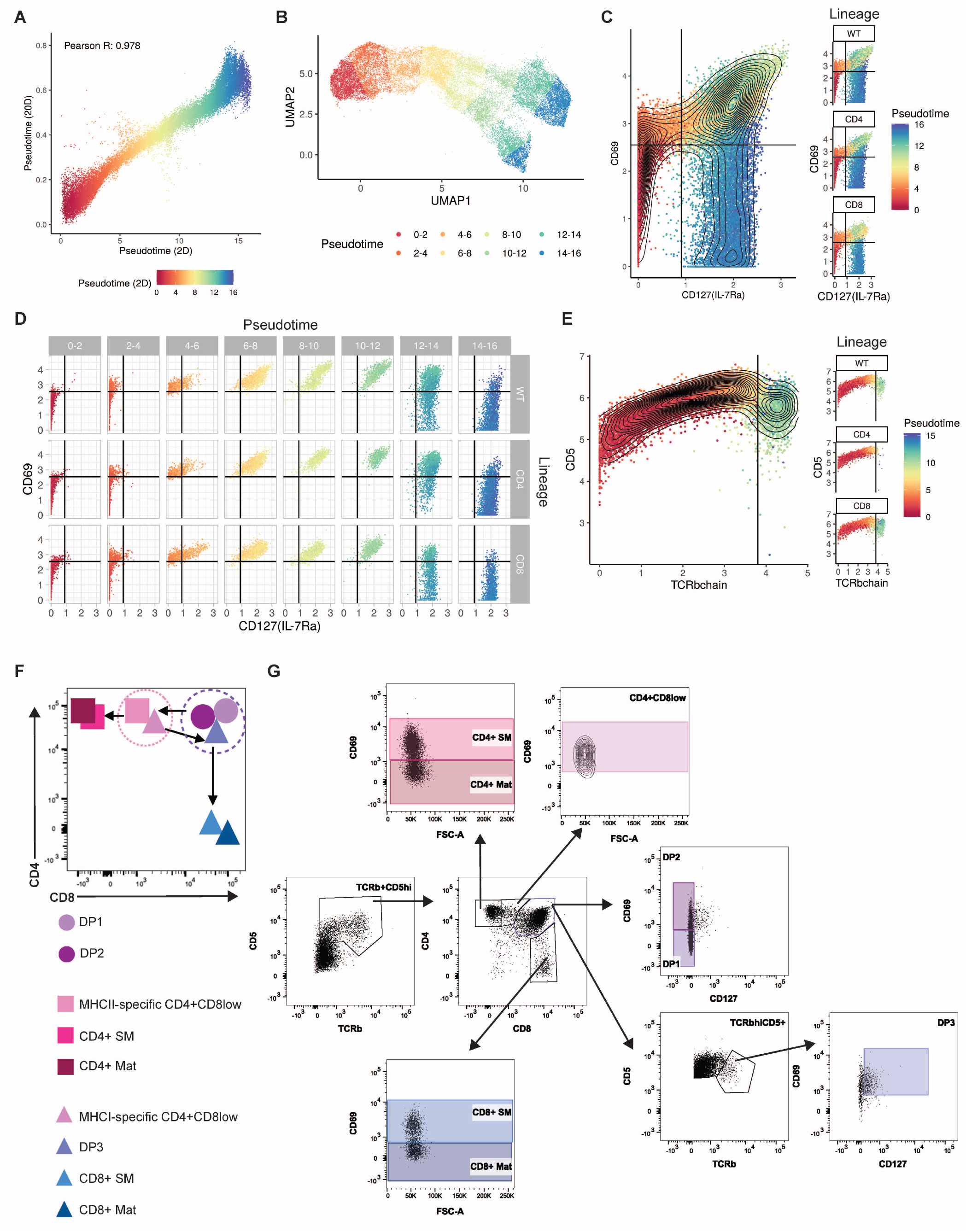
Pseudotime inference identifies intermediate thymocyte stages that can be isolated by FACS. **(A)** Correlation between Slingshot pseudotime inferred from the full 20-dimensional totalVI latent space and a 2-dimensional UMAP projection of the 20-dimensional latent space. **(B)** UMAP plot of the totalVI latent space from positively-selected thymocytes. Cells are colored according to placement in one of eight bins uniformly spaced over 2D pseudotime for visualization. **(C)** In silico flow cytometry plots of log(totalVI denoised expression) of CD127(IL-7Ra) and CD69 from positively-selected thymocytes (left) and the same cells separated by lineage (right). Cells are colored by pseudotime. **(D)** In silico flow cytometry plot of data as in (C) separated by lineage and pseudotime. **(E)** In silico flow cytometry plots of log(totalVI denoised expression) of TCRβ and CD5 from DP thymocytes (left) and the same cells separated by lineage (right). Cells are colored by pseudotime. Among DP thymocytes, the DP3 population is TCRβ high, CD127+, and CD69+. **(F)** Schematic of a CD4 versus CD8 biaxial plot to identify gated populations in adult thymocytes. Cells were gated into eight subsets: DP1, DP2, CD4+CD8low, semimature CD4 (CD4+ SM), mature CD4 (CD4+ Mat), DP3, semimature CD8 (CD8+ SM), and mature CD8 (CD8+ Mat). Circles represent lineage uncommitted cells, squares represent CD4 lineage cells, and triangles represent CD8 lineage cells. **(G)** Representative flow cytometry gating strategy for thymocyte populations in adult mice. Thymocytes were harvested from 6-8-week-old WT (C57BL/6 strain), MHCI-/- or MHCII-/- mice. Cell populations were gated to include lymphocytes, exclude forward scatter and side scatter doublets, include live cells, include TCRβ+CD5int/hi, then on CD4 versus CD8. Cell populations were gated into the following subsets based upon cell surface marker expression: DP1 (CD4+CD8+CD127-CD69-), DP2 (CD4+CD8+CD127-CD69+), CD4+CD8low (CD4+CD8low; CD4+CD8lowCD69+), DP3 (CD4+CD8+TCRβhiCD5+CD127+CD69+), semimature CD4 (CD4+ SM; CD4+CD8-CD69+), mature CD4 (CD4+ Mat; CD4+CD8-CD69-), semimature CD8 (CD8+ SM; CD8+CD69+), and mature CD8 (CD8+ Mat; CD8+CD69-).

**Figure S3:**
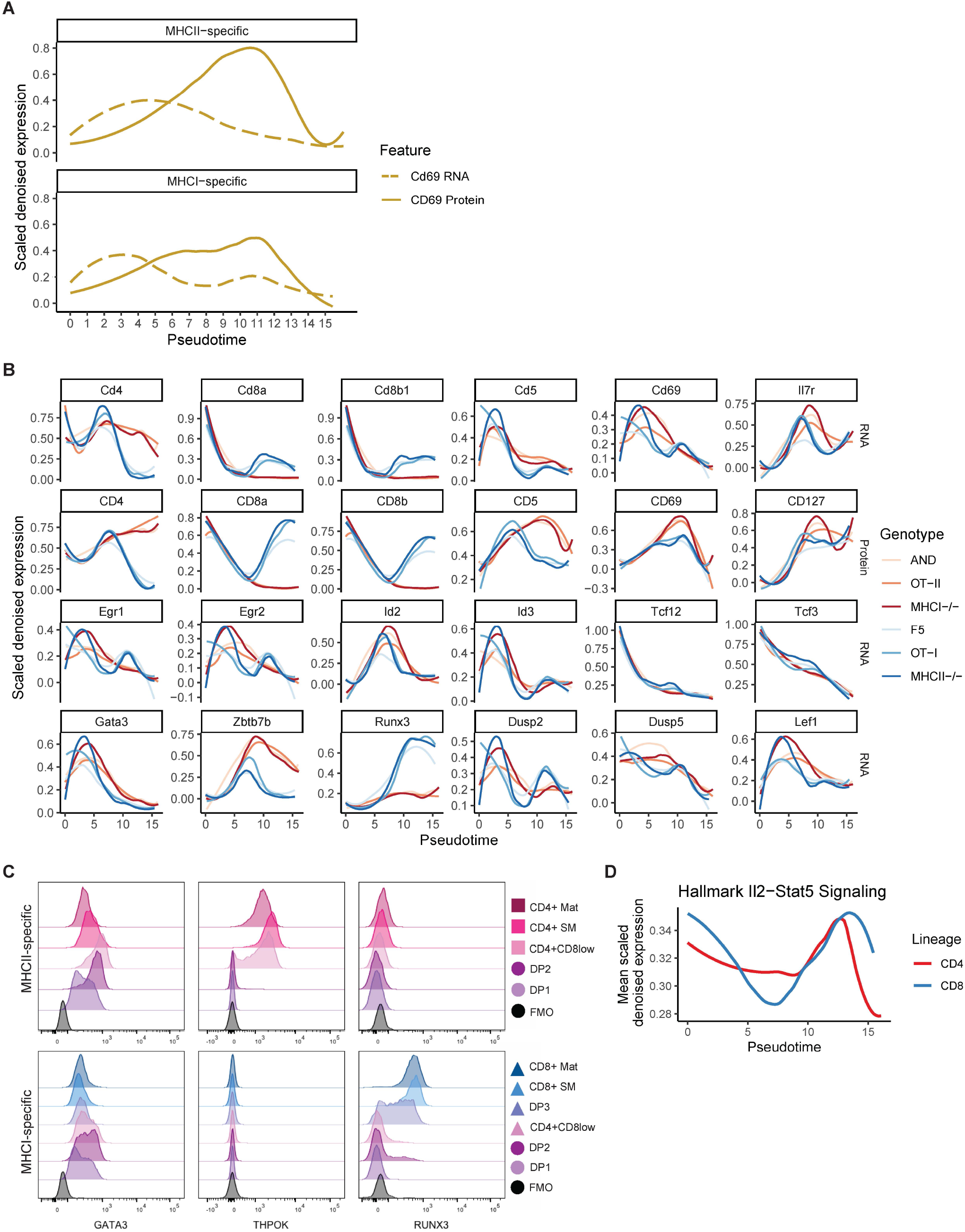
CITE-seq and fluorescence-based flow cytometry reveal the timing of expression for transcription factors and other features of CD4/CD8 development. **(A)** Expression over pseudotime of *Cd69* RNA (dashed) and CD69 protein (solid). Features are totalVI denoised expression values scaled per feature and smoothed by loess curves. **(B)** Expression of RNA and protein features over pseudotime by genotype. Features are totalVI denoised expression values scaled per feature and smoothed by loess curves. **(C)** Transcription factor protein expression in adult thymocyte populations. Representative histograms displaying GATA3, THPOK, and RUNX3 transcription factor expression detected by intracellular flow cytometry staining in MHCII-specific (MHCI-/-) and MHCI-specific (MHCII-/-) thymocyte populations. Thymocyte populations were gated on lymphocytes, excluding forward scatter and side scatter doublets, live cells, TCRβ+CD5int/hi then on CD4 versus CD8. Cell populations were gated into the following subsets based upon cell surface marker expression: DP1 (CD4+CD8+CD127-CD69-), DP2 (CD4+CD8+CD127-CD69+), DP3 (CD4+CD8+TCRβhiCD5+CD127+CD69+), CD4+CD8low (CD4+CD8lowCD69+), semimature CD4 (CD4+ SM; CD4+CD8-CD69+), mature CD4 (CD4+ Mat; CD4+CD8-CD69-), semimature CD8 (CD8+ SM; CD8+CD69+), and mature CD8 (CD8+ Mat; CD8+CD69-). Data is concatenated from n = 4 mice per genotype. Positive staining was determined using a fluorescence minus one (FMO) control. **(D)** Expression over pseudotime of the Hallmark Il2-Stat5 Signaling signature (Liberzon et al., 2011) displayed as the mean of scaled totalVI denoised expression per gene, smoothed by loess curves.

**Figure S4:**
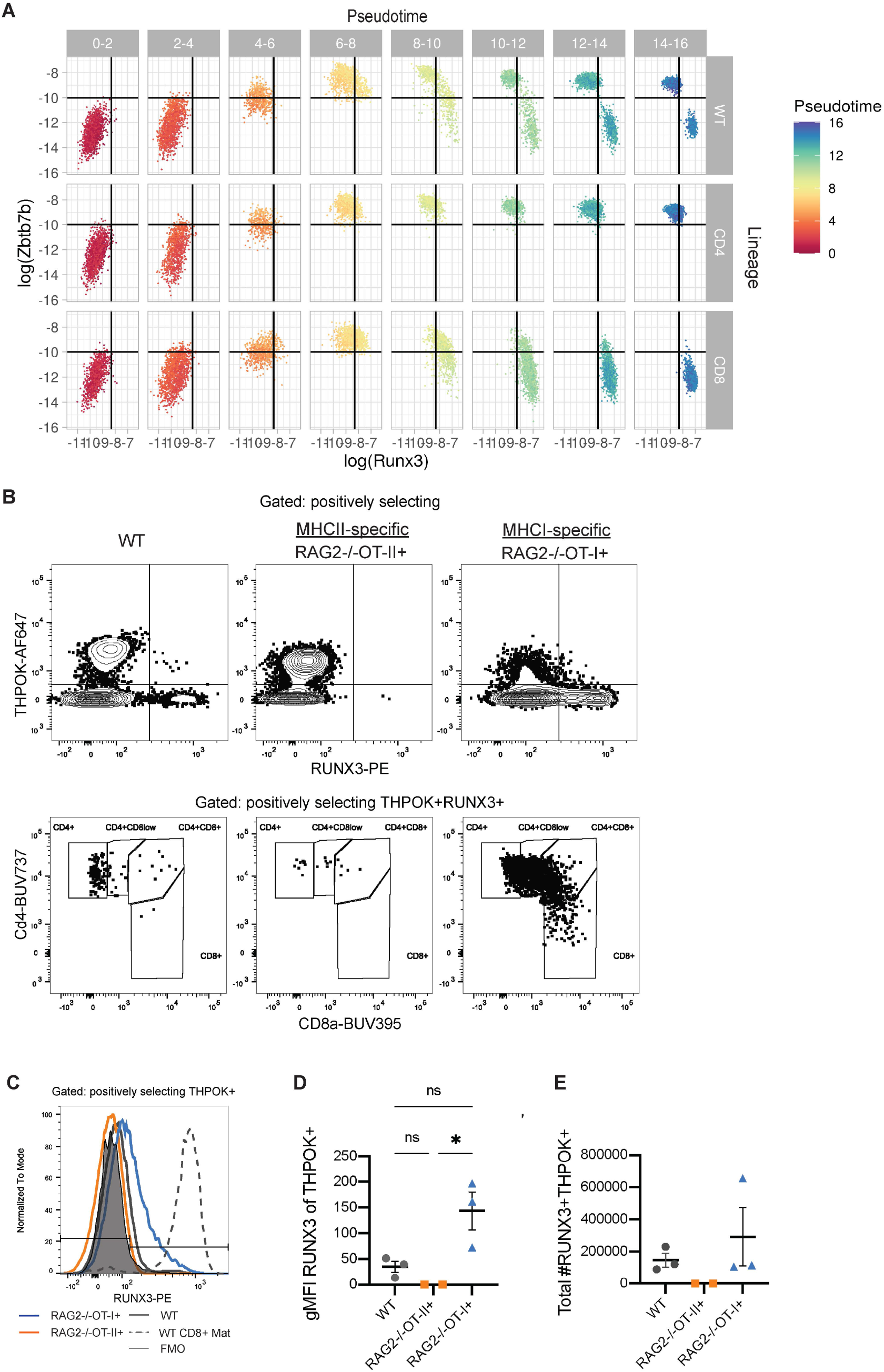
CITE-seq and fluorescence-based flow cytometry of key transcription factors. **(A)** In silico flow cytometry plots of log(totalVI denoised expression) of *Runx3* and *Zbtb7b* from positively-selected thymocytes separated by pseudotime. **(B)** Dual expression of THPOK and RUNX3 in positively selecting CD8-fated thymocytes. Top row shows representative flow cytometry contour plots of gated positively selecting thymocytes (CD5+, CD4+ or CD8+, CD24low/int) displaying RUNX3 vs THPOK protein expression from 6-8-week-old, adult WT, MHCII-specific TCR-tg RAG2-/-OT-II+, and MHCI-specific TCR-tg RAG2-/-OT-I+ mice. Positive staining and gates were determined using fluorescence minus one (FMO) controls. Bottom row shows representative FACS dot plots displaying CD8a vs CD4 expression in positively selecting RUNX3+THPOK+ thymocytes, **(C)** Representative histogram overlays displaying RUNX3 expression in THPOK+ positively selecting thymocytes from OT-II+/RAG2-/- (orange), OT-I+/RAG2-/- (blue) and WT (gray) mice. RUNX3 expression in CD8+ Mat cells (CD8+CD4-TCR+) from WT mice (gray, dashed line) is included for comparison. FMO is displayed as thin line, filled histogram (black). **(D)** Compiled data showing geometric mean fluorescent (gMFI) intensity of RUNX3 on THPOK+ positively selecting thymocytes. gMFI for each sample was calculated by subtracting the gMFI of the FMO. **(E)** Total number of positively selecting RUNX3+THPOK+ thymocytes in each mouse. Data is compiled from two independent experiments. In (D-E), each symbol represents 1 mouse. For WT mice (n=3), RAG2-/-OT-II+ (n=2), and RAG2-/-OT-I+ (n=3).

**Figure S5:**
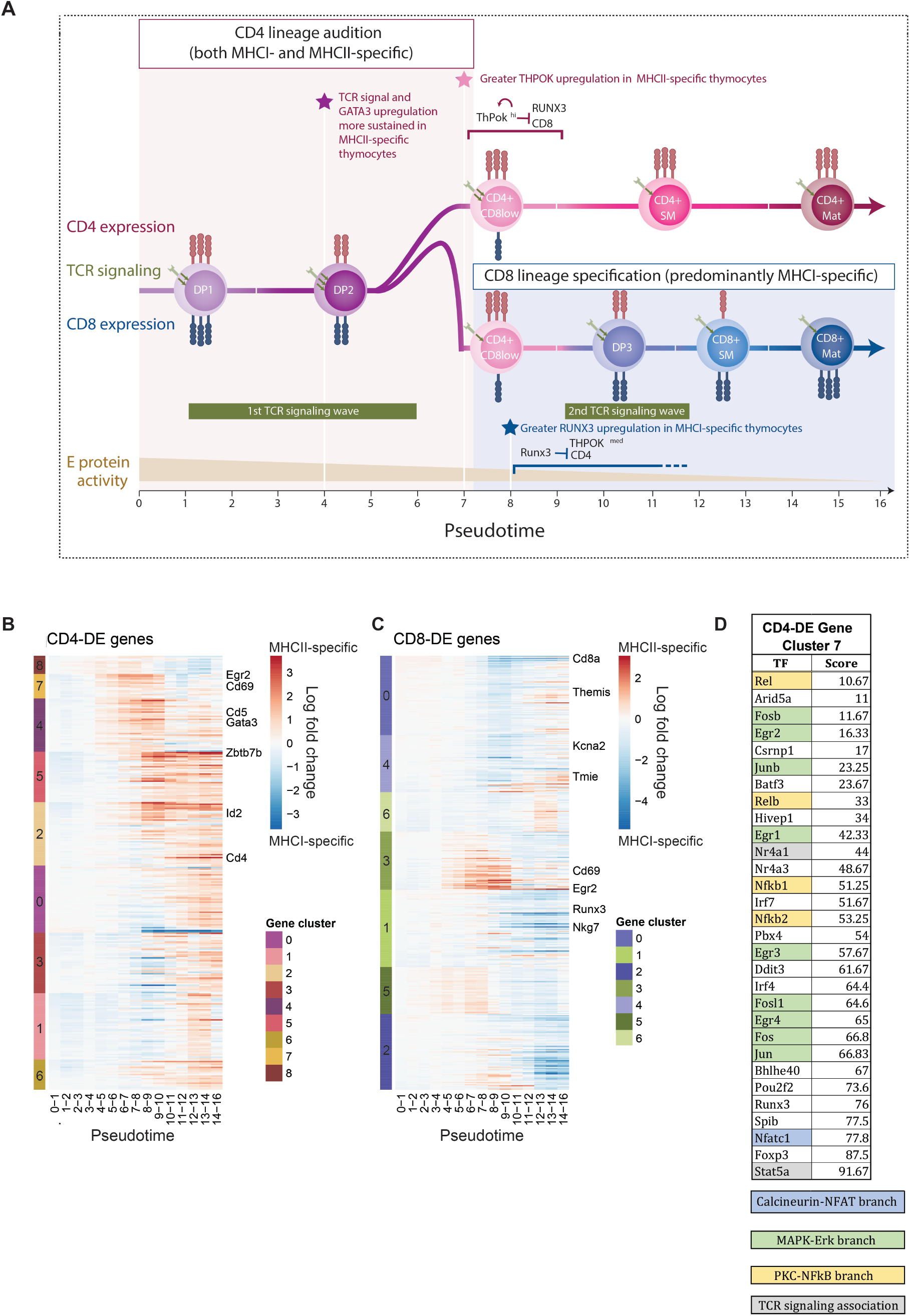
Model and differential expression analyses. **(A)** A sequential model for CD4 versus CD8 T cell lineage commitment. Key events during positive selection inferred from CITE-seq data are displayed from left to right in their order of occurrence based on pseudotime. Colored circles indicate the order of appearance of key thymocyte stages as defined by cell surface markers. Shaded red area indicates the time window during which both MHCI- and MHCII-specific thymocytes audition for the CD4 fate, corresponding to upregulation of GATA3 followed by THPOK. Shaded blue area indicates the later time window during which those thymocytes that failed the CD4 audition (mostly MHCI-specific) receive CD8 lineage reinforcement and survival signals. Green horizontal bars indicate two distinct temporal waves of TCR signaling: a first wave that is stronger and more sustained in MHCII-compared to MHCI-specific thymocytes, and a second later wave that occurs only in MHCI-specific thymocytes during the CD8 lineage specification phase. Stars indicate the key time points of lineage divergence, including the earliest detection of greater TCR signals and GATA3 upregulation in MHCII-specific thymocytes (purple star), followed by preferential THPOK induction and CD8 repression in MHCII-specific thymocytes (red star), and finally preferential RUNX3 induction and CD4 repression in MHCI-specific thymocytes (blue star). Red bracket indicates the time window during which MHCII-specific thymocytes commit to the CD4 lineage by fully upregulating THPOK, leading to activation of a THPOK autoregulation loop (Muroi et al., 2008) and full repression of CD8. Blue bracket indicates the time window during which MHCI-specific thymocytes turn on RUNX3, leading to repression of THPOK and CD4. Brown triangle indicates the continuous downregulation of E protein transcription factor activity throughout positive selection, which eventually allows for CD8 lineage specification in thymocytes that do not express high levels of THPOK (Jones-Mason et al., 2012). **(B)** totalVI median log fold change over pseudotime of genes upregulated in the CD4 lineage relative to the CD8 lineage. Genes are grouped by clusters shown in (Figure 4B). Clusters are ordered by their average highest magnitude fold change. **(C)** totalVI median log fold change over pseudotime of genes downregulated in the CD4 lineage relative to the CD8 lineage (i.e., upregulated in the CD8 lineage). Genes are grouped by clusters shown in (Figure 4C). Clusters are ordered by their average highest magnitude fold change. **(D)** Transcription factor (TF) enrichment analysis for TCR target-enriched gene clusters CD4-DE clusters 7. The top 30 TFs enriched in the gene set are shown. The full ChEA3 enrichment analysis is in Table S11 and S12. Colored boxes correspond to TFs activated by the respective branch of TCR signaling annotated in the inset diagram. Gray boxes indicate additional TFs associated with TCR signaling based on Netpath (Kandasamy et al., 2010), and as labeled in Figure 4E-F.

**Figure S6:**
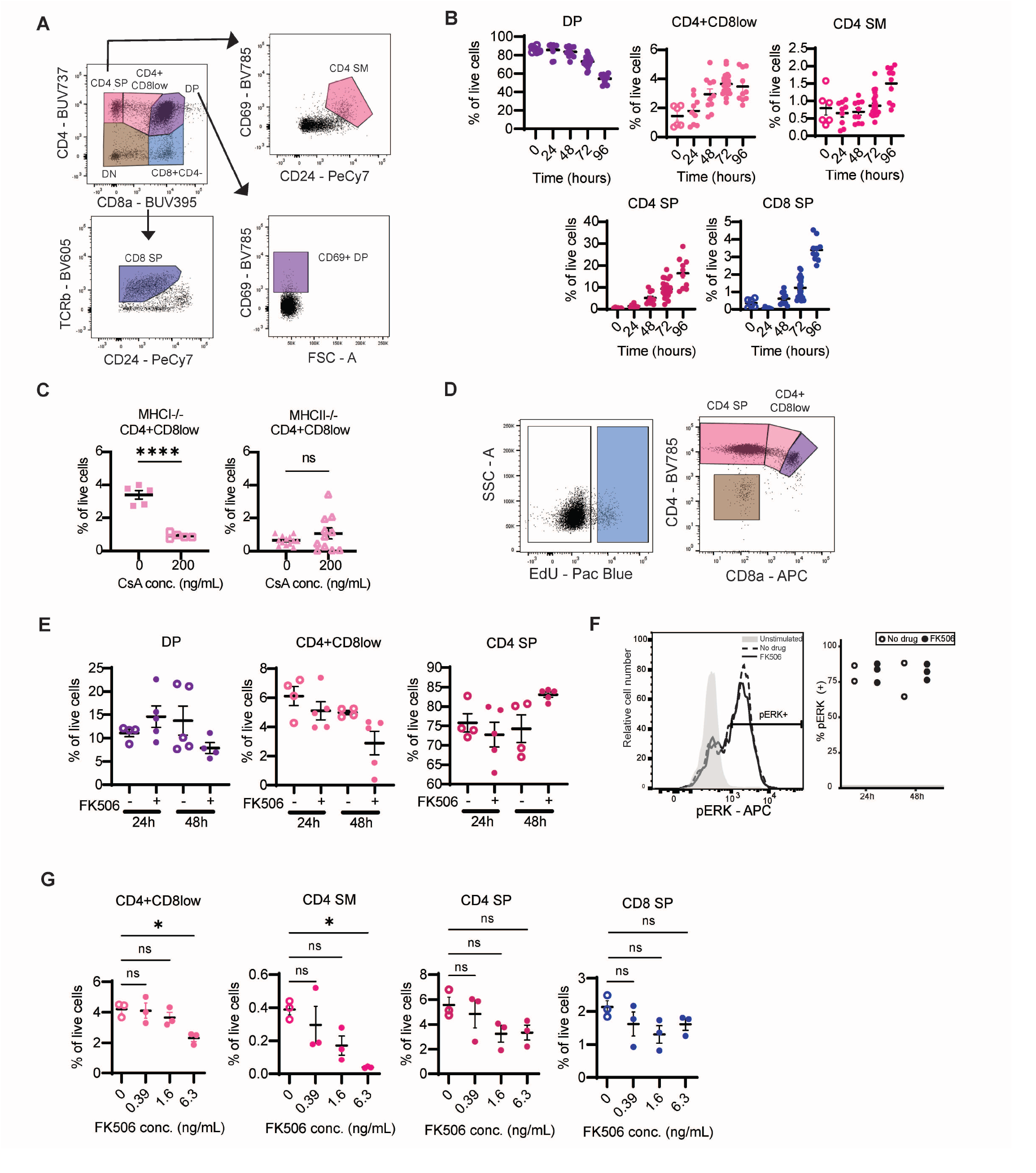
Inhibition of calcineurin blocks new CD4 SP development and GATA3 induction. **(A)** Representative flow cytometry gating strategy for neonatal thymic slice samples. **(B)** Time course of cell type proportion changes in neonatal slice cultures. Graphs display the frequency (% of live cells) of the indicated populations after 0, 24, 48, 72 and 96 hours of culture. Graphs contain data compiled from 9 independent experiments with WT slices. **(C)** Frequency (% of live cells) of CD4+CD8low cells in slices from MHCI-/- (MHCII-specific; squares) or MHCII-/- (MHCI-specific; triangles) mice following culture in medium alone (No CsA) or with 200 ng/mL CsA for 96 hours. Data are compiled from 2 independent experiments with MHCI-/- thymic slices and 5 independent experiments with MHCII-/- slices. **(D)** Gating strategy for in vivo EdU/FK506 experiment. **(E)** Frequency (% of live cells) of the indicated thymocyte populations with and without FK506. Each dot represents an individual mouse, and data are pooled from 3 (D) or 2 (E) independent experiments. **(F)** Thymocytes from FK506-treated or control AND TCRtg mice were stimulated via TCR crosslinking and analyzed by intracellular pERK staining and flow cytometry. Left panel shows a representative histogram of pERK induction in gated CD4+CD8+ thymocytes upon TCR crosslinking. Unstimulated samples (no crosslinking) are shown in gray. Right panel shows quantification of pERK induction (% of DP) with and without FK506 treatment. Data are compiled from 2 independent experiments, and each dot corresponds to a sample. **(G)** Frequency of the indicated populations after culture in the presence of the indicated concentration of FK506 for 72 hours. Each symbol on the graphs represents a thymic slice.

**Figure S7:**
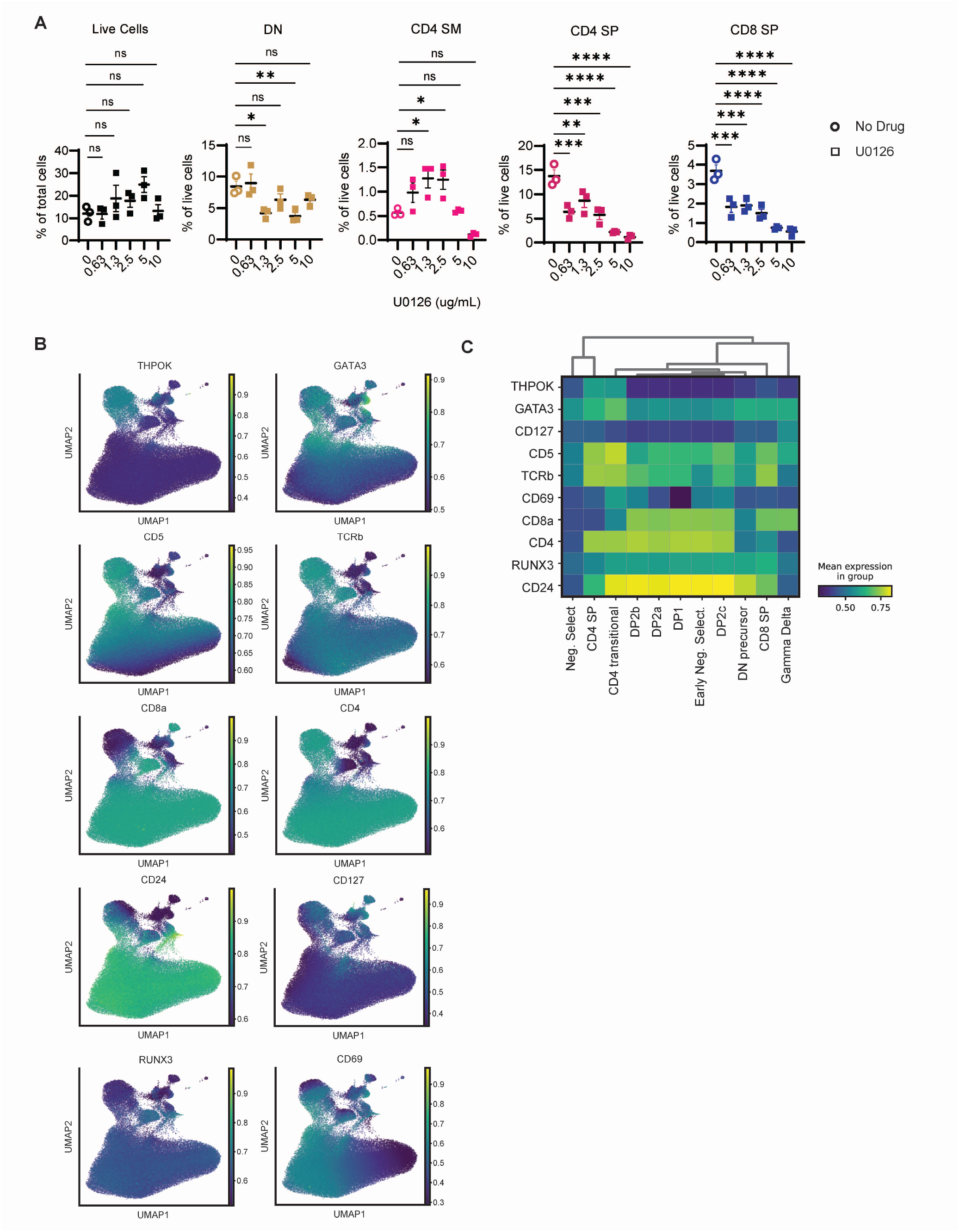
Multi-dimensional computation gating of neonatal slice cultures with calcineurin or MEK blockade. Thymic tissue slices were prepared from postnatal (day 1) mice and cultured with either calcineurin inhibitor CsA or MEK inhibitor U0126. **(A)** U0126-treated thymic slices were incubated in concentrations ranging from 10 μg/mL to 0.63 μg/mL. CD4 SM, CD4 SP and CD8 SP, plots show frequency (%) out of live cells. Plots are representative of 3 independent experiments. **(B)** UMAP plots of multi-dimensional flow cytometry data colored by scaled expression of each flow cytometry marker. Data are pooled samples from the representative experiment shown in Figures 6A-C and S8. **(C)** Heatmap of relative protein expression of markers in each subset shown in Figures 6A-C and S8.

**Figure S8:**
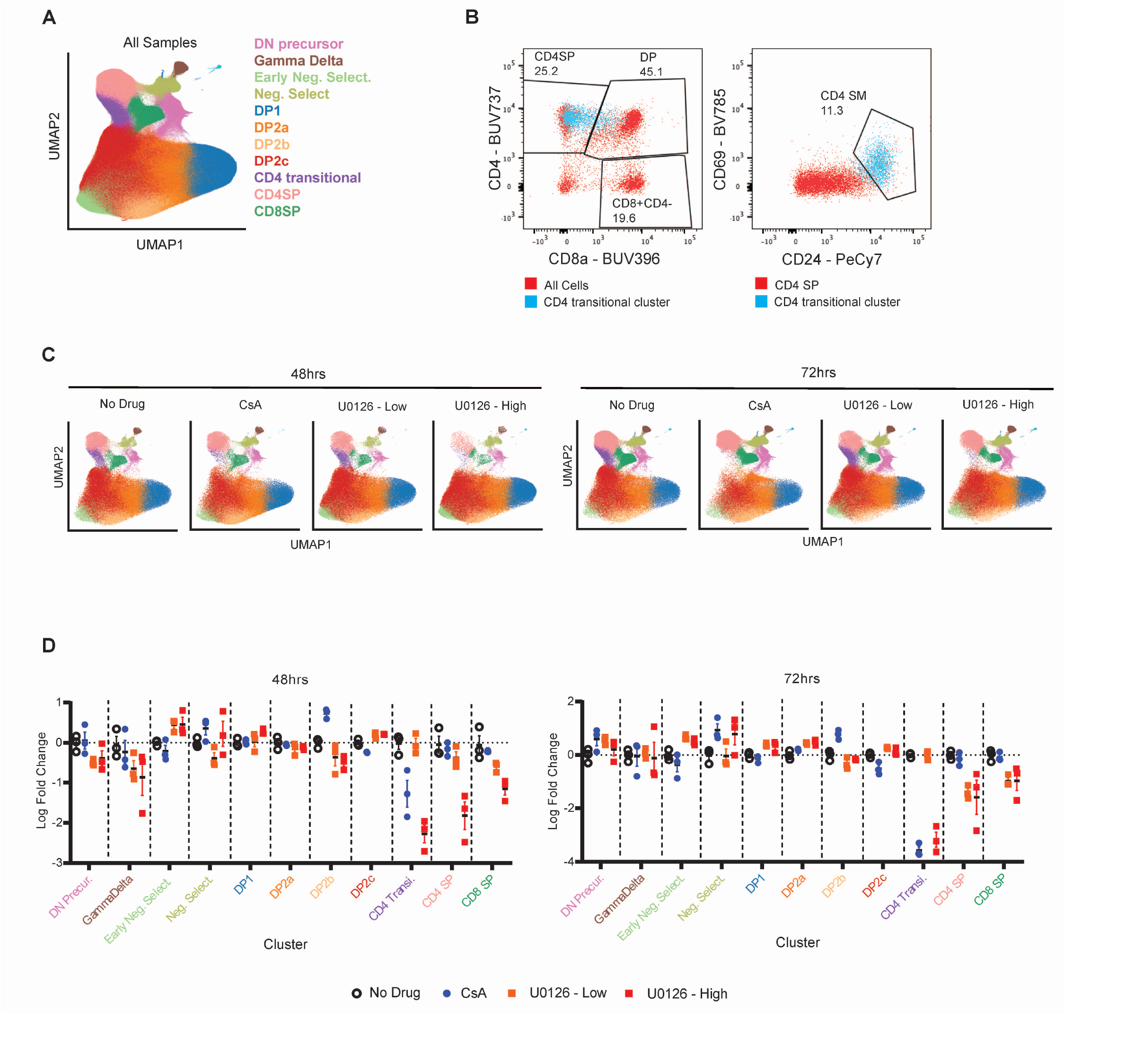
Calcineurin inhibition selectively impacts the CD4 audition. Thymic tissue slices were prepared from postnatal (day 1) mice and cultured with either calcineurin inhibitor CsA (200 ng/mL) or MEK inhibitor U0126 (2 μg/mL or 10 μg/mL). Thymic slices were collected at 48 or 72 hours, stained with fluorescent antibodies and analyzed by either manual or computational, multidimensional gating. **(A)** UMAP based on multi-dimensional flow cytometry data of all samples. **(B)** Cells from the CD4 transitional cluster defined by computational gating (blue) are superimposed on live gated cells (red, left hand plot) or CD4+CD8-gated cells (red, right hand plot) for comparison with manual gating strategy. A 48 hour, no drug control sample is shown. **(C)** UMAP plot by the indicated experimental condition. **(D)** Scatter plots showing the log fold change in cell type proportion relative to no drug control for indicated cell clusters for each condition, separated by time (left panel is 48 hours; right panel is 72 hours). Data is from one representative experiment out of two.

## References

Au-Yeung, B. B., Melichar, H. J., Ross, J. O., Cheng, D. A., Zikherman, J., Shokat, K. M., … Weiss, A. (2014). Quantitative and temporal requirements revealed for Zap70 catalytic activity during T cell development. Nature Immunology, 15(7), 687–694. https://doi.org/10.1038/ni.2918

Becht, E., McInnes, L., Healy, J., Dutertre, C.-A., Kwok, I. W. H., Ng, L. G., … Newell, E. W. (2019). Dimensionality reduction for visualizing single-cell data using UMAP. Nature Biotechnology, 37(1), 38–44. https://doi.org/10.1038/nbt.4314

Bhakta, N. R., Oh, D. Y., & Lewis, R. S. (2005). Calcium oscillations regulate thymocyte motility during positive selection in the three-dimensional thymic environment. Nature Immunology, 6(2), 143–151. https://doi.org/10.1038/ni1161

Bosselut, R. (2004). CD4/CD8-lineage differentiation in the thymus: From nuclear effectors to membrane signals. Nature Reviews Immunology. Nature Publishing Group. https://doi.org/10.1038/nri1392

Cannarile, M. A., Lind, N. A., Rivera, R., Sheridan, A. D., Camfield, K. A., Wu, B. B., … Goldrath, A. W. (2006). Transcriptional regulator Id2 mediates CD8+ T cell immunity. Nature Immunology, 7(12), 1317–1325. https://doi.org/10.1038/ni1403

Chakraborty, A. K., & Weiss, A. (2014). Insights into the initiation of TCR signaling. Nature Immunology. Nature Publishing Group. https://doi.org/10.1038/ni.2940

Chan, S. H., Cosgrove, D., Waltzinger, C., Benoist, C., & Mathis, D. (1993). Another view of the selective model of thymocyte selection. Cell, 73(2), 225–236. https://doi.org/10.1016/0092-8674(93)90225-F

Chan, S., Waltzinger, C., Tarakhovsky, A., Benoist, C., & Mathis, D. (1999). An influence of CD5 on the selection of CD4-lineage T cells. European Journal of Immunology, 29(9), 2916–2922. https://doi.org/10.1002/(SICI)1521-4141(199909)29:09<2916::AID-IMMU2916>3.0.CO;2-I

Chèneby, J., Ménétrier, Z., Mestdagh, M., Rosnet, T., Douida, A., Rhalloussi, W., … Ballester, B. (2020). ReMap 2020: A database of regulatory regions from an integrative analysis of Human and Arabidopsis DNA-binding sequencing experiments. Nucleic Acids Research, 48(D1), D180–D188. https://doi.org/10.1093/nar/gkz945

Choi, S., Cornall, R., Lesourne, R., & Love, P. E. (2017, September 1). THEMIS: Two Models, Different Thresholds. Trends in Immunology. Elsevier Ltd. https://doi.org/10.1016/j.it.2017.06.006

Chopp, L. B., Gopalan, V., Ciucci, T., Ruchinskas, A., Rae, Z., Lagarde, M., … Bosselut, R. (2020). An Integrated Epigenomic and Transcriptomic Map of Mouse and Human ab T Cell Development. Immunity, 53(6). https://doi.org/10.1016/j.immuni.2020.10.024

Chung, H., Parkhurst, C. N., Magee, E. M., Phillips, D., Habibi, E., Chen, F., … Regev, A. (2021). Simultaneous single cell measurements of intranuclear proteins and gene expression. BioRxiv, 2021.01.18.427139. https://doi.org/10.1101/2021.01.18.427139

Daley, S. R., Hu, D. Y., & Goodnow, C. C. (2013). Helios marks strongly autoreactive CD4+ T cells in two major waves of thymic deletion distinguished by induction of PD-1 or NF-κB. Journal of Experimental Medicine, 210(2), 269–285. https://doi.org/10.1084/jem.20121458

d’Ambrosio, D., Cantrell, D. A., Frati, L., Santoni, A., & Testi, R. (1994). Involvement of p21ras activation in T cell CD69 expression. European Journal of Immunology, 24(3), 616–620. https://doi.org/10.1002/eji.1830240319

Daniels, M. A., Teixeiro, E., Gill, J., Hausmann, B., Roubaty, D., Holmberg, K., … Palmer, E. (2006). Thymic selection threshold defined by compartmentalization of Ras/MAPK signalling. Nature, 444(7120), 724–729. https://doi.org/10.1038/nature05269

DeTomaso, D., Jones, M. G., Subramaniam, M., Ashuach, T., Ye, C. J., & Yosef, N. (2019). Functional interpretation of single cell similarity maps. Nature Communications, 10(1). https://doi.org/10.1038/s41467-019-12235-0

Dobin, A., Davis, C. A., Schlesinger, F., Drenkow, J., Zaleski, C., Jha, S., … Gingeras, T. R. (2013). STAR: ultrafast universal RNA-seq aligner. Bioinformatics, 29(1), 15–21. https://doi.org/10.1093/bioinformatics/bts635

Donnadieu, E., Lang, V., Bismuth, G., Ellmeier, W., Acuto, O., Michel, F., & Trautmann, A. (2001). Differential Roles of Lck and Itk in T Cell Response to Antigen Recognition Revealed by Calcium Imaging and Electron Microscopy. The Journal of Immunology, 166(9), 5540–5549. https://doi.org/10.4049/jimmunol.166.9.5540

Dunham, I., Kundaje, A., Aldred, S. F., Collins, P. J., Davis, C. A., Doyle, F., … Lochovsky, L. (2012). An integrated encyclopedia of DNA elements in the human genome. Nature, 489(7414), 57–74. https://doi.org/10.1038/nature11247

Dzhagalov, I. L., Melichar, H. J., Ross, J. O., Herzmark, P., & Robey, E. A. (2012). Two-Photon Imaging of the Immune System. In Current Protocols in Cytometry (Vol. 60, pp. 12.26.1-12.26.20). Hoboken, NJ, USA: John Wiley & Sons, Inc. https://doi.org/10.1002/0471142956.cy1226s60

Egawa, T., & Littman, D. R. (2008). ThPOK acts late in specification of the helper T cell lineage and suppresses Runx-mediated commitment to the cytotoxic T cell lineage. Nature Immunology, 9(10), 1131–1139. https://doi.org/10.1038/ni.1652

Gallo, E. M., Winslow, M. M., Canté-Barrett, K., Radermacher, A. N., Ho, L., McGinnis, L., … Crabtree, G. R. (2007). Calcineurin sets the bandwidth for discrimination of signals during thymocyte development. Nature, 450(7170), 731–735. https://doi.org/10.1038/nature06305

Gayoso, A., Steier, Z., Lopez, R., Regier, J., Nazor, K. L., Streets, A., & Yosef, N. (2021). Joint probabilistic modeling of single-cell multi-omic data with totalVI. Nature Methods, 18(3). https://doi.org/10.1038/s41592-020-01050-x

Germain, R. N. (2002). t-cell development and the CD4-CD8 lineage decision. Nature Reviews Immunology. European Association for Cardio-Thoracic Surgery. https://doi.org/10.1038/nri798

Germain, R. N., Robey, E. A., & Cahalan, M. D. (2012, June 29). A Decade of imaging cellular motility and interaction dynamics in the immune system. Science. American Association for the Advancement of Science. https://doi.org/10.1126/science.1221063

Gimferrer, I., Hu, T., Simmons, A., Wang, C., Souabni, A., Busslinger, M., … Alberola-Ila, J. (2011). Regulation of GATA-3 Expression during CD4 Lineage Differentiation. The Journal of Immunology, 186, 3892–3898. https://doi.org/10.4049/jimmunol.1003505

Grusby, M. J., Johnson, R. S., Papaioannou, V. E., & Glimcher, L. H. (1991). Depletion of CD4+ T cells in major histocompatibility complex class II-deficient mice. Science, 253(5026), 1417–1420. https://doi.org/10.1126/science.1910207

Hedrick, S. M., Michelini, R. H., Doedens, A. L., Goldrath, A. W., & Stone, E. L. (2012, September). FOXO transcription factors throughout T cell biology. Nature Reviews Immunology. NIH Public Access. https://doi.org/10.1038/nri3278

Hettmann, T., & Leiden, J. M. (2000). NF-κB Is Required for the Positive Selection of CD8 + Thymocytes. The Journal of Immunology, 165(9), 5004–5010. https://doi.org/10.4049/jimmunol.165.9.5004

Hogquist, K. A., & Jameson, S. C. (2014). The self-obsession of T cells: How TCR signaling thresholds affect fate “decisions” and effector function. Nature Immunology, 15(9), 815–823. https://doi.org/10.1038/ni.2938

Hogquist, K., Xing, Y., Hsu, F.-C., & Shapiro, V. S. (2015). T Cell Adolescence: Maturation Events Beyond Positive Selection. Journal of Immunology, 195(4), 1351–1357. https://doi.org/10.4049/jimmunol.1501050

Hu, Q., Nicol, S. A., Suen, A. Y. W., & Baldwin, T. A. (2012). Examination of thymic positive and negative selection by flow cytometry. Journal of Visualized Experiments, (68). https://doi.org/10.3791/4269

Itano, A., Salmon, P., Kioussis, D., Tolaini, M., Corbella, P., & Robey, E. (1996). The cytoplasmic domain of CD4 promotes the development of CD4 lineage T cells. Journal of Experimental Medicine, 183(3), 731–741. https://doi.org/10.1084/jem.183.3.731

Jimi, E., Strickland, I., Voll, R. E., Long, M., & Ghosh, S. (2008). Differential Role of the Transcription Factor NF-κB in Selection and Survival of CD4+ and CD8+ Thymocytes. Immunity, 29(4), 523–537. https://doi.org/10.1016/j.immuni.2008.08.010

Jones-Mason, M. E., Zhao, X., Kappes, D., Lasorella, A., Iavarone, A., & Zhuang, Y. (2012). E Protein Transcription Factors Are Required for the Development of CD4 + Lineage T Cells. Immunity, 36(3), 348–361. https://doi.org/10.1016/j.immuni.2012.02.010

Kandasamy, K., Sujatha Mohan, S., Raju, R., Keerthikumar, S., Sameer Kumar, G. S., Venugopal, A. K., … Pandey, A. (2010). NetPath: A public resource of curated signal transduction pathways. Genome Biology, 11(1), R3. https://doi.org/10.1186/gb-2010-11-1-r3

Karimi, M. M., Guo, Y., Cui, X., Pallikonda, H. A., Horková, V., Wang, Y. F., … Merkenschlager, M. (2021). The order and logic of CD4 versus CD8 lineage choice and differentiation in mouse thymus. Nature Communications, 12(1), 1–14. https://doi.org/10.1038/s41467-020-20306-w

Kaye, J., Hsu, M. L., Sauron, M. E., Jameson, S. C., Gascoigne, N. R. J., & Hedrick, S. M. (1989). Selective development of CD4+ T cells in transgenic mice expressing a class II MHC-restricted antigen receptor. Nature, 341(6244), 746–749. https://doi.org/10.1038/341746a0

Keenan, A. B., Torre, D., Lachmann, A., Leong, A. K., Wojciechowicz, M. L., Utti, V., … Ma’ayan, A. (2019). ChEA3: transcription factor enrichment analysis by orthogonal omics integration. Nucleic Acids Research, 47(W1), W212–W224. https://doi.org/10.1093/nar/gkz446

Kernfeld, E. M., Genga, R. M. J., Neherin, K., Magaletta, M. E., Xu, P., & Maehr, R. (2018). A Single-Cell Transcriptomic Atlas of Thymus Organogenesis Resolves Cell Types and Developmental Maturation. Immunity, 48(6), 1258–1270.e6. https://doi.org/10.1016/j.immuni.2018.04.015

Kisielow, P., & Miazek, A. (1995). Positive selection of T cells: Rescue from programmed cell death and differentiation require continual engagement of the T cell receptor. Journal of Experimental Medicine, 181(6), 1975–1984. https://doi.org/10.1084/jem.181.6.1975

Kovanen, P. E., Bernard, J., Al-Shami, A., Liu, C., Bollenbacher-Reilley, J., Young, L., Pise-Masison, C., Spolski, R., & Leonard, W. J. (2008). T-cell development and function are modulated by dual specificity phosphatase DUSP5. The Journal of Biological Chemistry, 283(25), 17362–17369. https://doi.org/10.1074/jbc.M70988720

Kurd, N., & Robey, E. A. (2016). T-cell selection in the thymus: A spatial and temporal perspective. Immunological Reviews, 271(1), 114–126. https://doi.org/10.1111/imr.12398

Lavaert, M., Liang, K. L., Vandamme, N., Park, J.-E., Roels, J., Kowalczyk, M. S., … Taghon, T. (2020). Integrated scRNA-Seq Identifies Human Postnatal Thymus Seeding Progenitors and Regulatory Dynamics of Differentiating Immature Thymocytes. Immunity, 52(6). https://doi.org/10.1016/j.immuni.2020.03.019

Liu, J. (1993). FK506 and cyclosporin, molecular probes for studying intracellular signal transduction. Immunology Today, 14(6), 290–295. https://doi.org/10.1016/0167-5699(93)90048-P

Liu, X., & Bosselut, R. (2004). Duration of TCR signaling controls CD4-CD8 lineage differentiation in vivo. Nature Immunology, 5(3), 280–288. https://doi.org/10.1038/ni1040

Lopez, R., Regier, J., Cole, M. B., Jordan, M. I., & Yosef, N. (2018). Deep generative modeling for single-cell transcriptomics. Nature Methods, 15(12), 1053–1058. https://doi.org/10.1038/s41592-018-0229-2

López-Rodríguez, C., Aramburu, J., & Berga-Bolaños, R. (2015). Transcription factors and target genes of pre-TCR signaling. Cellular and Molecular Life Sciences, 72(12), 2305–2321. https://doi.org/10.1007/s00018-015-1864-8

Lucas, B., & Germain, R. N. (1996). Unexpectedly complex regulation of CD4/CD8 coreceptor expression supports a revised model for CD4+CD8+ thymocyte differentiation. Immunity, 5(5), 461–477. https://doi.org/10.1016/S1074-7613(00)80502-6

Lucas, B., Vasseur, F., & Penit, C. (1993). Normal Sequence of Phenotypic Transitions in One Cohort of 5=Bromo=2’-Deoxyuridine-Pulse-Labeled Thymocytes Correlation with T Cell Receptor Expression’ (Vol. 151). Retrieved from http://www.jimmunol.org/

Liberzon, A., Subramanian, A., Pinchbakc, R., Thorvaldsdottir, H., Tamayo, P., Mesirov, J. (2011). Molecular signatures database (MSigDB) 3.0. Bioinformatics, 27(12), 1739–1740. https://doi.org/10.1093/bioinformatics/btr260

Lundberg, K., Heath, W., Köntgen, F., Carbone, F. R., & Shortman, K. (1995). Intermediate steps in positive selection: Differentiation of CD4+8int TCRint thymocytes into CD4−8+TCRhi Thymocytes. Journal of Experimental Medicine, 181(5), 1643–1651. https://doi.org/10.1084/jem.181.5.1643

Lutes, L. K., Steier, Z., McIntyre, L. L., Pandey, S., Kaminski, J., Hoover, A. R., … Robey, E. A. (2021). T cell self-reactivity during thymic development dictates the timing of positive selection. ELife, 10. https://doi.org/10.7554/eLife.65435

Malissen, B., Grégoire, C., Malissen, M., & Roncagalli, R. (2014). Integrative biology of T cell activation. Nature Immunology. Nature Publishing Group. https://doi.org/10.1038/ni.2959

Mamalaki, C., Norton, T., Tanaka, Y., Townsend, A. R., Chandler, P., Simpson, E., & Kioussis, D. (1992). Thymic depletion and peripheral activation of class I major histocompatibility complex-restricted T cells by soluble peptide in T-cell receptor transgenic mice. Proceedings of the National Academy of Sciences of the United States of America, 89(23), 11342–11346. https://doi.org/10.1073/pnas.89.23.11342

Marodon, G., & Rocha, B. (1994). Generation of mature T cell populations in the thymus: CD4 or CD8 down-regulation occurs at different stages of thymocyte differentiation. European Journal of Immunology, 24(1), 196–204. https://doi.org/10.1002/eji.1830240131

Matechak, E. O., Killeen, N., Hedrick, S. M., & Fowlkes, B. J. (1996). MHC class II-specific T cells can develop in the CD8 lineage when CD4 is absent. Immunity, 4(4), 337–347. https://doi.org/10.1016/S1074-7613(00)80247-2

McNeil, L. K., Starr, T. K., & Hogquist, K. A. (2005). A requirement for sustained ERK signaling during thymocyte positive selection in vivo. Proceedings of the National Academy of Sciences of the United States of America, 102(38), 13574–13579. https://doi.org/10.1073/pnas.0505110102

Melichar, H. J., Ross, J. O., Herzmark, P., Hogquist, K. A., & Robey, E. A. (2013). Distinct Temporal Patterns of T Cell Receptor Signaling During Positive Versus Negative Selection in Situ. Science Signaling, 6(297), ra92–ra92. https://doi.org/10.1126/scisignal.2004400

Mingueneau, M., Jiang, W., Feuerer, M., Mathis, D., & Benoist, C. (2012). Thymic negative selection is functional in NOD mice. Journal of Experimental Medicine, 209(3), 623–637. https://doi.org/10.1084/jem.20112593

Mingueneau, M., Kreslavsky, T., Gray, D., Heng, T., Cruse, R., Ericson, J., … Turley, S. (2013). The transcriptional landscape of αβ T cell differentiation. Nature Immunology, 14(6), 619–632. https://doi.org/10.1038/ni.2590

Moran, A. E., Holzapfel, K. L., Xing, Y., Cunningham, N. R., Maltzman, J. S., Punt, J., & Hogquist, K. A. (2011). T cell receptor signal strength in Treg and iNKT cell development demonstrated by a novel fluorescent reporter mouse. Journal of Experimental Medicine, 208(6), 1279–1289. https://doi.org/10.1084/jem.20110308

Muroi, S., Naoe, Y., Miyamoto, C., Akiyama, K., Ikawa, T., Masuda, K., … Taniuchi, I. (2008). Cascading suppression of transcriptional silencers by ThPOK seals helper T cell fate. Nature Immunology, 9(10), 1113–1121. https://doi.org/10.1038/ni.1650

Nabet, B., Roberts, J. M., Buckley, D. L., Paulk, J., Dastjerdi, S., Yang, A., Leggett, A. L., Erb, M. A., Lawlor, M. A., Souza, A., Scott, T. G., Vittori, S., Perry, J. A., Qi, J., Winter, G. E., Wong, K.-K., Gray, N. S., & Bradner, J. E. (2018). The dTAG system for immediate and target-specific protein degradation. Nature Chemical Biology, 14(5), 431–441. https://doi.org/10.1038/s41589-018-0021-8

Navarro, M. N., & Cantrell, D. A. (2014, August 19). Serine-threonine kinases in TCR signaling. Nature Immunology. Nature Publishing Group. https://doi.org/10.1038/ni.2941

Park, J. E., Botting, R. A., Conde, C. D., Popescu, D. M., Lavaert, M., Kunz, D. J., … Teichmann, S. A. (2020). A cell atlas of human thymic development defines T cell repertoire formation. Science, 367(6480). https://doi.org/10.1126/science.aay3224

Perera, E. M., Bao, Y., Kos, L., & Berkovitz, G. (2010). Structural and functional characterization of the mouse tescalcin promoter. Gene, 464(1–2), 50–62. https://doi.org/10.1016/J.GENE.2010.06.002

Ross, J. O., Melichar, H. J., Au-Yeung, B. B., Herzmark, P., Weiss, A., & Robey, E. A. (2014). Distinct phases in the positive selection of CD8+ T cells distinguished by intrathymic migration and T-cell receptor signaling patterns. Proceedings of the National Academy of Sciences of the United States of America, 111(25). https://doi.org/10.1073/pnas.1408482111

Ross, J. O., Melichar, H. J., Halkias, J., & Robey, E. A. (2015). Studying T cell development in thymic slices. In T-Cell Development: Methods and Protocols (Vol. 1323, pp. 131–140). Springer New York. https://doi.org/10.1007/978-1-4939-2809-5_11

Saelens, W., Cannoodt, R., Todorov, H., & Saeys, Y. (2019). A comparison of single-cell trajectory inference methods. Nature Biotechnology, 37(5), 547–554. https://doi.org/10.1038/s41587-019-0071-9

Saini, M., Sinclair, C., Marshall, D., Tolaini, M., Sakaguchi, S., & Seddon, B. (2010). Regulation of Zap70 expression during thymocyte development enables temporal separation of CD4 and CD8 repertoire selection at different signaling thresholds. Science Signaling, 3(114), ra23–ra23. https://doi.org/10.1126/scisignal.2000702

Scheinman, E. J., & Avni, O. (2009). Transcriptional regulation of Gata3 in T helper cells by the integrated activities of transcription factors downstream of the interleukin-4 receptor and T cell receptor. Journal of Biological Chemistry, 284(5), 3037–3048. https://doi.org/10.1074/jbc.M807302200

Seong, R. H., Chamberlain, J. W., & Parnes, J. R. (1992). Signal for T-cell differentiation to a CD4 cell lineage is delivered by CD4 transmembrane region and/or cytoplasmic tail. Nature, 356(6371), 718–720. https://doi.org/10.1038/356718a0

Shao, H., Kono, D. H., Chen, L. Y., Rubin, E. M., & Kaye, J. (1997). Induction of the early growth response (Egr) family of transcription factors during thymic selection. Journal of Experimental Medicine, 185(4), 731–744. https://doi.org/10.1084/jem.185.4.731

Sharp, L. L., Schwarz, D. A., Bott, C. M., Marshall, C. J., & Hedrick, S. M. (1997). The influence of the MAPK pathway on T cell lineage commitment. Immunity, 7(5), 609–618. https://doi.org/10.1016/S1074-7613(00)80382-9

Shinzawa, M., Moseman, E. A., Gossa, S., Mano, Y., Bhattacharya, A., Guinter, T., Alag, A., Chen, X., Cam, M., McGavern, D. B., Erman, B., & Singer, A. (2022). Reversal of the T cell immune system reveals the molecular basis for T cell lineage fate determination in the thymus. Nature Immunology. https://doi.org/10.1038/s41590-022-01187-1

Sinclair, C., & Seddon, B. (2014). Overlapping and Asymmetric Functions of TCR Signaling during Thymic Selection of CD4 and CD8 Lineages. The Journal of Immunology, 192(11), 5151–5159. https://doi.org/10.4049/jimmunol.1303085

Singer, A., Adoro, S., & Park, J. H. (2008, October). Lineage fate and intense debate: Myths, models and mechanisms of CD4-versus CD8-lineage choice. Nature Reviews Immunology. Nat Rev Immunol. https://doi.org/10.1038/nri2416

Stassen, S.V., Siu, D.M.D., Lee, K.C.M., Ho, J.W.K., So, H.K.H., Tsia, K.K. (2020). PARC: ultrafast and accurate clustering of phenotypic data of millions of single cells. Bioinformatics, 36(9), 2778–2786. http://doi.org/10.1093/bioinformatics/btaa042.

Stoeckius, M., Hafemeister, C., Stephenson, W., Houck-Loomis, B., Chattopadhyay, P. K., Swerdlow, H., … Smibert, P. (2017). Simultaneous epitope and transcriptome measurement in single cells. Nature Methods, 14(9), 865–868. https://doi.org/10.1038/nmeth.4380

Street, K., Risso, D., Fletcher, R. B., Das, D., Ngai, J., Yosef, N., … Dudoit, S. (2018). Slingshot: Cell lineage and pseudotime inference for single-cell transcriptomics. BMC Genomics, 19(1), 1–16. https://doi.org/10.1186/s12864-018-4772-0

Stuart, T., Butler, A., Hoffman, P., Hafemeister, C., Papalexi, E., Mauck, W. M., … Satija, R. (2019). Comprehensive Integration of Single-Cell Data. Cell, 177(7), 1888–1902.e21. https://doi.org/10.1016/j.cell.2019.05.031

Taniuchi, I. (2016). Views on helper/cytotoxic lineage choice from a bottom-up approach. Immunological Reviews, 271(1), 98–113. https://doi.org/10.1111/imr.12401

Tanzola, M. B., & Kersh, G. J. (2006). The dual specificity phosphatase transcriptome of the murine thymus. Molecular Immunology, 43(6), 754–762. https://doi.org/10.1016/j.molimm.2005.03.006

Traag, V. A., Waltman, L., & van Eck, N. J. (2019). From Louvain to Leiden: guaranteeing well-connected communities. Scientific Reports, 9(1), 1–12. https://doi.org/10.1038/s41598-019-41695-z

Vacchio, M. S., & Bosselut, R. (2016). Function Transcriptional Circuitry To Control Their Lineage Commitment + CD8 − + in the Thymus: How T Cells Recycle the CD4 What Happens in the Thymus Does Not Stay. J Immunol References, 196, 4848–4856. https://doi.org/10.4049/jimmunol.1600415

Wang, C. R., Hashimoto, K., Kubo, S., Yokochi, T., Kubo, M., Suzuki, M., Suzuki, K., Tada, T., & Nakayama, T. (1995). T cell receptor-mediated signaling events in CD4+CD8+ thymocytes undergoing thymic selection: requirement of calcineurin activation for thymic positive selection but not negative selection. The Journal of Experimental Medicine, 181(3), 927–941. https://doi.org/10.1084/jem.181.3.927

Wang, L., Wildt, K. F., Zhu, J., Zhang, X., Feigenbaum, L., Tessarollo, L., … Bosselut, R. (2008). Distinct functions for the transcription factors GATA-3 and ThPOK during intrathymic differentiation of CD4+ T cells. Nature Immunology, 9(10), 1122–1130. https://doi.org/10.1038/ni.1647

Wang, L., Xiong, Y., & Bosselut, R. (2010, October 1). Tenuous paths in unexplored territory: From T cell receptor signaling to effector gene expression during thymocyte selection. Seminars in Immunology. Academic Press. https://doi.org/10.1016/j.smim.2010.04.013

Weist, B. M., Kurd, N., Boussier, J., Chan, S. W., & Robey, E. A. (2015). Thymic regulatory T cell niche size is dictated by limiting IL-2 from antigen-bearing dendritic cells and feedback competition. Nature Immunology, 16(6), 635–641. https://doi.org/10.1038/ni.3171

Wilkinson, B., & Kaye, J. (2001). Requirement for sustained MAPK signaling in both CD4 and CD8 lineage commitment: A threshold model. Cellular Immunology, 211(2), 86–95. https://doi.org/10.1006/cimm.2001.1827

Wolf, F. A., Angerer, P., & Theis, F. J. (2018). SCANPY: Large-scale single-cell gene expression data analysis. Genome Biology. https://doi.org/10.1186/s13059-017-1382-0

Wolf, F.A., Hamey, F.K., Plass, M. et al. (2019). PAGA: graph abstraction reconciles clustering with trajectory inference through a topology preserving map of single cells. Genome Biology, 20(59), https://doi.org/10.1186/s13059-019-1663-x

Wong, W. F., Looi, C. Y., Kon, S., Movahed, E., Funaki, T., Chang, L. Y., … Kohu, K. (2014). T-cell receptor signaling induces proximal Runx1 transactivation via a calcineurin-NFAT pathway. European Journal of Immunology, 44(3), 894–904. https://doi.org/10.1002/eji.201343496

Xing, Y., Wang, X., Jameson, S. C., & Hogquist, K. A. (2016). Late stages of T cell maturation in the thymus involve NF-κB and tonic type i interferon signaling. Nature Immunology, 17(5), 565–573. https://doi.org/10.1038/ni.3419

Xiong, Y., & Bosselut, R. (2012, April 1). CD4-CD8 differentiation in the thymus: Connecting circuits and building memories. Current Opinion in Immunology. Elsevier Current Trends. https://doi.org/10.1016/j.coi.2012.02.002

Yan, Y., Zhang, G. X., Williams, M. S., Carey, G. B., Li, H., Yang, J., … Xu, H. (2012). TCR stimulation upregulates MS4a4B expression through induction of AP-1 transcription factor during T cell activation. Molecular Immunology, 52(2), 71–78. https://doi.org/10.1016/j.molimm.2012.04.011

Yasutomo, K., Doyle, C., Miele, L., & Germain, R. N. (2000). The duration of antigen receptor signalling determines CD4+ versus CD8+ T-cell lineage fate. Nature, 404(6777), 506–510. https://doi.org/10.1038/35006664

Zamisch, M., Tian, L., Grenningloh, R., Xiong, Y., Wildt, K. F., Ehlers, M., … Bosselut, R. (2009). The transcription factor Ets1 is important for CD4 repression and Runx3 up-regulation during CD8 T cell differentiation in the thymus. Journal of Experimental Medicine, 206(12), 2685–2699. https://doi.org/10.1084/jem.20092024

Zheng, G. X. Y., Terry, J. M., Belgrader, P., Ryvkin, P., Bent, Z. W., Wilson, R., … Bielas, J. H. (2017). Massively parallel digital transcriptional profiling of single cells. Nature Communications, 8, 14049. https://doi.org/10.1038/ncomms14049

Zhou, W., Yui, M. A., Williams, B. A., Yun, J., Wold, B. J., Cai, L., & Rothenberg, E. V. (2019). Single-Cell Analysis Reveals Regulatory Gene Expression Dynamics Leading to Lineage Commitment in Early T Cell Development. Cell Systems, 9(4), 321–337.e9. https://doi.org/10.1016/j.cels.2019.09.008

